# Generative modeling of short, disordered proteins with homogeneous sequence composition

**DOI:** 10.1101/2022.11.11.516154

**Authors:** Ishan Taneja, Keren Lasker

## Abstract

Protein design has seen remarkable progress in the past decade, with numerous examples of *de novo* proteins with novel topologies and functions being successfully synthesized. Computational tools have played a large role in the ability to rationally design proteins. Recently, there have been numerous successes applying deep learning techniques to protein design that have demonstrated comparable or significantly improved performance over traditional energy-based approaches. However, the protein design problem has been less well explored for disordered proteins or, more generally, proteins with conformational heterogeneity. In this work, we demonstrate that if one approximates the spatial output of a coarse-grained molecular dynamics simulation as a multivariate normal distribution parameterized by a mean vector (representing an ensemble-averaged pairwise distance map) and covariance matrix, one can train a generative model to learn the distribution of these parameters across a set of sequences. Specifically, we encoded the mean vector and covariance matrix for each sequence in a low-dimensional space via a fixed linear transformation and trained a masked auto-encoder to accurately learn the distribution of this low-dimensional output. Furthermore, by sampling from the masked auto-encoder and transforming the generated samples back into their original high-dimensional space, one can generate realistic, ensemble-averaged pairwise distance maps. These results were demonstrated on coarse-grained simulation data derived from approximately 2000 distinct sequences, each sequence being 24 residues in length and consisting exclusively of glycine, serine, glutamate, and lysine. Though this set of sequences is relatively homogeneous in composition, we speculate our approach can be applied to disordered sequences of longer length and more heterogeneous composition, given the appropriate training set.

## 1 Introduction

Computational tools have played a large role in the ability to rationally design proteins [1, 2]. Recently, there have been numerous successes applying deep learning techniques to protein design that have demonstrated comparable or significantly improved performance over traditional energy based approaches [3, 4, 5, 6]. To date, protein design efforts have primarily focused on designing sequences with a specific tertiary structure. A much less well explored question is how to design disordered proteins with certain properties or functions. This problem is not as clearly defined as designing folded proteins, as disordered proteins are highly dynamic and tend to have a broad free energy landscape rather than a global free energy minima [7, 8, 9, 10]. Furthermore, evidence suggests that properties of disordered proteins may be more of a function of local and global sequence features as opposed to the precise sequence itself [11, 12, 13, 14, 15]. Thus, there could be a large set of sequences with heterogeneous composition that are disordered and satisfy a target property. This contrasts with the structured protein design problem, where a relatively limited set of sequences adopt a specific target fold.

One key aspect to the disordered protein design problem is specifying the target property of interest. We note that the ideal target property would contain high spatial and temporal resolution (e.g all-atom < *x, y, z* > coordinates sampled at the nanosecond timescale). This is because the ability to design sequences with a high-dimensional target property implies the ability to design sequences with a low-dimensional target property, but not vice-versa.

Given the complex and potentially ambiguous nature of the disordered protein design problem, we elected to focus on a simpler but related question. Specifically, given high spatial resolution information (*d*) for a set of disordered sequences, can we train a model to accurately learn *p*(*d*)? We believe the ability to do this would serve as a useful stepping stone to tackling the more ambitious problem of learning *p*(*s*|*d*) (i.e the distribution of sequences with high-dimensional target property *d*).

Because experimental techniques such as small-angle X-ray scattering (SAXS) or single-molecule fluorescence resonance energy transfer (smFRET) generally report ensemble averaged structural parameters such as the radius of gyration (*R_g_*) or end-to-end distance [16], we elected to use molecular dynamic simulations to generate a dataset with high spatial and temporal resolution information for a large set of disordered sequences. We specifically used the recently developed Mpipi model [17], which has demonstrated state of the art accuracy on the ability to predict single and multi-chain properties. To simplify the learning problem, we exclusively trained and evaluated the model on data generated for 24 residue long sequences consisting exclusively of glycine (G), serine (S), glutamate (E), and lysine (K). At a high level, our approach consists of two stages: (i) generating a low-dimensional representation of the data (*x*) via a fixed, invertible linear transformation, and (ii) training a masked auto-encoder to learn *p*(*x*). Our results demonstrate that 1) it is possible to accurately approximate *d* from *x*, 2) the model is able to learn an accurate representation of *p*(*x*) and 3) the model can generate realistic, averaged pairwise distance maps when compared to that of real disordered sequences. We suggest this approach serves as a promising way to learn *p*(*d*) for disordered sequences of longer length and more heterogeneous composition.

## 2 Methods

### 2.1 Coarse-grained molecular dynamic simulations

For a given sequence, canonical-ensemble simulations were performed at 300 K for 10 μs, with a time step of 10 fs. A Langevin thermostat was used, with a relaxation time of 50 fs. Trajectory frames were outputted every 1 ns, yielding approximately 3e4 frames per sequence. Three repeats were run per unique sequence.

For our force field, we used the Mpipi model, which implicitly represents each amino acid by a single bead. Briefly, the potential energy of a protein is modeled as the sum of bond energy, electrostatic energy, and a non-bonded Lennard-Jones like pair potential. The electrostatic term is represented as a screened coulomb interaction modeled by the Debye-Huckel approximation. The debye screening length was set to 795 pm, corresponding to a monovalent salt concentration of .15 M.

### 2.2 Sequences for which simulations were run

There were three sets of sequences, each 24 residues long, for which the simulations were run. In the first set, we restricted the sequence composition while in the second and third set, no explicit restrictions were placed. The first set of sequences were used to train and quantitatively evaluate the machine learning model, while the second and third set of sequences were used to inspect the limitations of the model on sequences with greater residue diversity.

The first set (Set 1) of sequences consisted exclusively of glycine (G), serine (S), glutamate (E), and lysine (K), with the constraint that 1/3 of the residues were either positively or negatively charged. We elected to constrain this set of sequences to these residues to simplify the learning problem. This first set of sequences can be divided into two classes. The first class of sequences are of the form GS*GS*GS*GS*GS*GS*GS*GS* (referred to as (GS*)_8_), where * refers to either E or K. We enumerated all combinations of sequences of this form (28 total). The second class of sequences took each of these 256 sequences, and generated permutations of the sequence such that its charge patterning parameter (*κ*) was equal to either {.1,.2,.3,….,.9}. This class of sequences is referred to as the *κ* permutations. *κ* is a parameter ranging between 0 and 1 that describes the extent of charged amino acid mixing in a sequence [18]. Low values of *κ* correspond to sequences that are well mixed with respect to its oppositely charged residues while higher values correspond to sequences with a higher degree of segregation between oppositely charged residues. For reference, these permuted sequences were generated using the python tool GOOSE [19] (https://github.com/idptools/goose). This procedure yielded a maximum of 10 permutations for each of the 256 sequences (certain sequence/*κ* constraints were unable to yield a sequence). In total, 1696 unique, permuted sequences were generated.

The second set (Set 2) of sequences (referred to as *κ* variants) was generated using functionality in GOOSE that enables one to design a disordered sequence with no restriction on sequence composition with an arbitrary value of *κ*. We generated 30 sequences per *κ* value in the set {.1, .3, .5, .7}. This yielded a total of 120 sequences. We only accepted a sequence in this set if its hydrophobicity was between 2.7 and 3.0, given that the hydrophobicity of all sequences in the first class were within that range. The third set (Set 3) of sequences (referred to as hydrophobic variants) was generated using the standard functionality in GOOSE to design an arbitrary disordered sequence. We only accepted a sequence in this set if its hydrophobicity was greater than 3.0, yielding 120 total sequences.

Detailed information regarding sequence properties for sequences in each of the three sets is provided in Fig. 1 and Fig. S1-3 [20].

**Figure 1:**
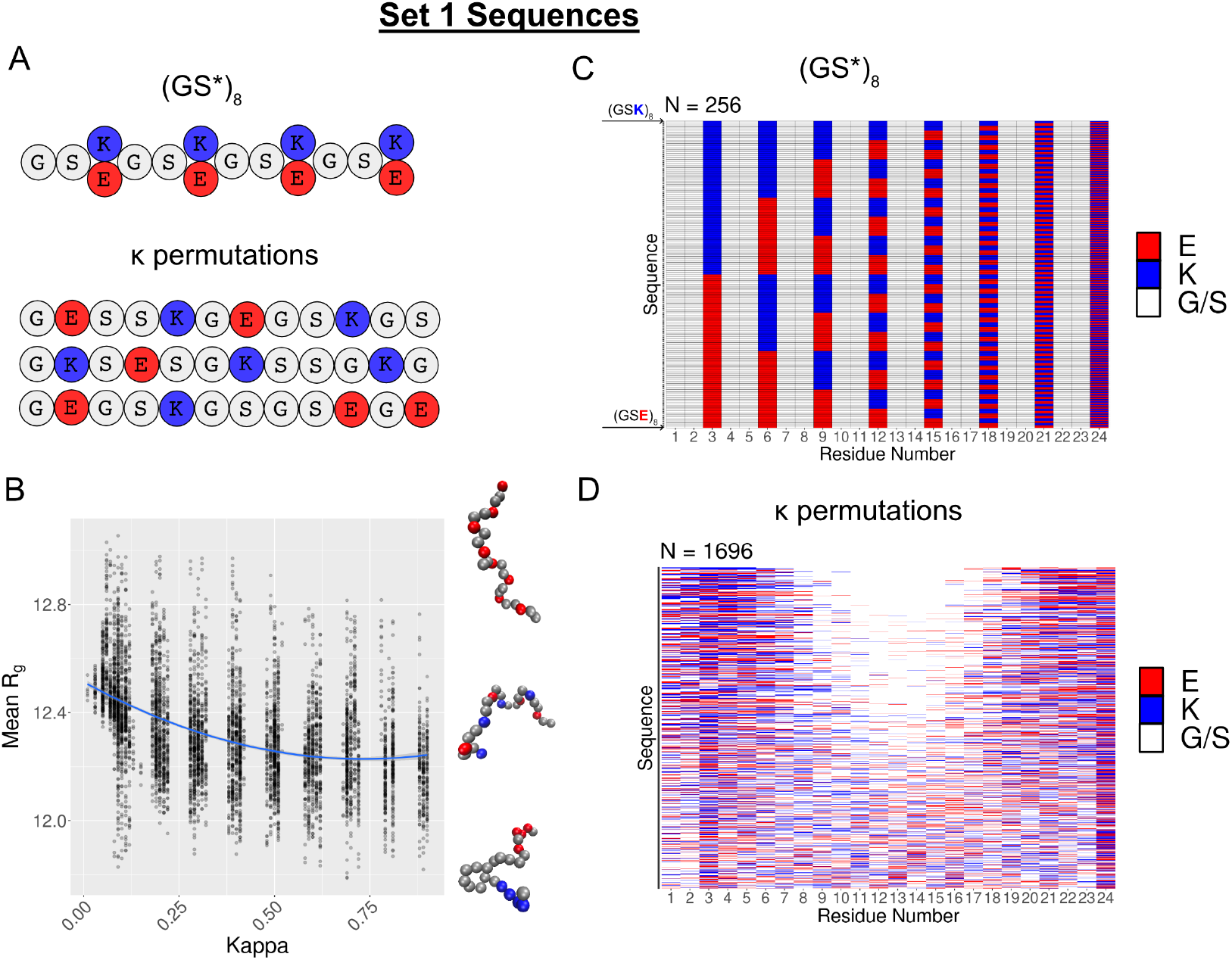
Detailed sequence information for (GS*)_8_ and *κ* permutations. A) The first class of sequences are of the form (GS*)_8_, where * refers to either E or K. We enumerated all combinations of sequences of this form (28 total). The second class of sequences took each of these 256 sequences, and generated permutations of the sequence such that *κ* was equal to either [.1, .2, .3, …, .9]. B) Relationship between the mean radius of gyration (*R_g_*) and *κ* across all sequences. Each point corresponds to a unique sequence in the (GS*)_8_ and *κ* permutations set. A second order polynomial was fit to the data for reference. The spearman correlation between *κ* and *R_g_* is −.46. C/D) The y-axis corresponds to a given sequence in either the (GS*)_8_ or *κ* permutations class. For each sequence, its residue is color coded according to its identity (Glutamate = Red, Lysine = Blue, Glycine/Serine = White)

### 2.3 Representing the output of a coarse-grained polymer simulation in a low-dimensional space

Coarse-grained molecular dynamics simulations provide < *x, y, z* >_*t*_ backbone coordinate information for each amino acid as a function of time. While the true data generating distribution of this process is known (i.e the Boltzmann distribution of form *e*^-*u*(*x*)^ where *x* is a microscopic configuration defined by backbone coordinates and *u* is the simulation specific potential energy function), there is no closed-form analytical representation of it. At the expense of losing temporal information, a middle ground approach is to solely represent the pairwise distance distribution of the coordinate data matrix with a multivariate Gaussian. Assuming a protein comprising *n* residues, the mean vector *μ* is an *n*^2^-dimensional vector where each element represents the mean distance between each pair of residues while Σ is an *n*^2^ by *n*^2^ matrix where each element represents the covariance between each pair of pairwise distances. For reference, we will refer to **M** as the *n* by *n* matrix representation of *μ*, where each element represents the mean distance between each pair of residues. In addition, we will refer to Σ as **C** to avoid ambiguity with notation later introduced.

The parameters defining this distribution are still relatively high-dimensional, requiring on the order *O*(*n*^2^ + *n*^4^) parameters. Given this, we sought to find a lower-dimensional representation of the data. Our approach is based on the observation that the underlying structure of **M** is shared across all proteins in our simulation (Fig. S4 and S5). Specifically, **E**[*d*(*i, j*)] ∝ |*i* – *j*|, where *i* and *j* represent an arbitrary pair of residues and **E**[*d*(*i, j*)] represents their average distance. This observation motivated us to learn the underlying fingerprint of each pairwise distance distribution, as opposed to extraneous features shared across sequences. To do so, we attempted to reconstruct each **M** and **C** using a common basis. Specifically, we calculated the singular value decomposition of **M**_*GS*_ and **C**_*GS*_ for a 24 residue polymer consisting of 12 consecutive glycine serine repeats (referred to as (GS)_12_). Both **M** and **C** can be decomposed as 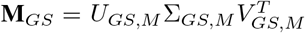 and 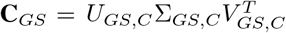, where Σ_*GS, M*_ and Σ_*GS, C*_ are diagonal matrices with non-negative diagonal entries that represent the singular values and *U/V* represent the left/right singular vectors. Given this, for an arbitrary sequence *s*, we can calculate an approximation of the singular values of **M**_s_/**C**_s_ with respect to the (GS)_12_ basis as follows: if we let 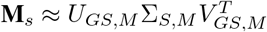 and impose the constraint that Σ_*S, M*_ is a diagonal matrix, then 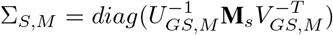.

We then tested whether **M** and **C** for an arbitrary sequence could each be reconstructed with tolerable errors (Fig. S6-S10). Mathematically, the reconstruction error of **M**/**C** is quantified as the spectral norm of 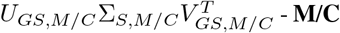. Our analyses suggested 1) the matrices could be faithfully reconstructed using the (GS)_12_ basis and 2) the reconstruction error levels off relatively quickly, suggesting only a subset of the singular values may be relevant to learn.

### 2.4 Learning the singular value distribution using a conditional masked autoencoder

From the simulation data for each sequence in our dataset, we can generate Σ_*S, M*_ and Σ_*S, C*_ according to the procedure defined in the preceding paragraph. As opposed to generating a single Σ_*S, M*_ and Σ_*S, C*_ per sequence, we generated 100 Σ_*S, M*_ and Σ_*S, C*_ per sequence, where each Σ_*S, M*_ and Σ_*S, C*_ is generated from 1000 frames randomly sampled with replacement among the cumulative 3*e*4 frames. We observed that the singular value distribution from these subsamples remained roughly identical to that of the one derived from the entire simulation (Fig. S11 and S12).

For reference, Σ_*S, M*_ consists of n values across its diagonal while Σ_*S, C*_ consists of *n*^2^ values diagonal. As opposed to learning all values in Σ_*S, C*_ and Σ_*S, G*_, we elected to learn the first eight diagonal entries of each matrix. Recent work has demonstrated that masked autoencoders are powerful models to estimate distributions. We employed a conditional version of this model, where instead of learning *p*(*x*), we learned *p*(*x*|*c*). In this case, *c* represents two covariates derived from the sequence: the net charge per residue (ncpr) and charge patterning parameter (*κ*). Each layer in the model is a feedforward neural network with masked weight matrices, such that the autoregressive property holds.

The procedure for designing the masks follows the standard approach of Germain et al [21]. Each input node is assigned a degree, which is an integer ranging from [0, *D_x_* + *D_c_* – 1]. *D_x_* represents the dimensionality of *x* (16: 8 singular values originating from Σ_*S,M*_ and 8 singular values originating from Σ_*S, C*_) while *D_c_* represents the dimensionality of the conditional covariates (2: ncpr and *κ*). Each hidden unit node is also assigned a degree, which is an integer ranging from [0, *D_x_* – 2]. Finally, each output unit node is assigned a degree, which is an integer ranging from [0, *D_x_* – 1]. The degree of an input node is taken to be its index in the order < 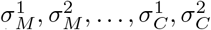, …, ncpr, *κ* > where 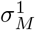 refers to the upper left diagonal entry of Σ_*S, M*_ and 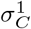 refers to the upper left diagonal entry of Σ_*S, C*_. The degree of an output node is taken to be its index in the order 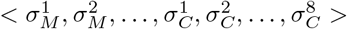. In order to enforce the autoregressive property, a hidden unit is allowed to receive input only from units with a lower or equal degree and an output unit is allowed to receive input only from units with a lower degree. Finally, for each input in *x*, there are two corresponding output nodes representing a prediction of its mean and standard deviation.

The model employed two hidden layers, each with 100 nodes. All models were trained with the Adam optimizer [22], using a minibatch size of 100, a step size of 10^−3^, and for 20 consecutive epochs. We employed a Gaussian negative log likelihood loss for the loss function.

### 2.5 Sampling from the conditional masked autoencoder

In order to sample from the model, we first generated a prediction of 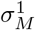 given *c*. Specifically, given our training set, we fit a second-order polynomial, *f*_2_, modeling 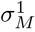 with *c* as the input. For a given *c*, we calculated the 90% prediction interval of *f*_2_(*c*) and sampled a random value from this interval. While theoretically we could sample 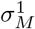 from the masked autoencoder itself by inputting < 0,0,0…, ncpr, *κ* >, the model was unable to learn a meaningful relationship between these variables. We speculate that the non-continuous nature of the ncpr and lack of smoothness of the distribution of *κ* may be contributing factors (Fig. S1). After generating 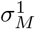, we generated 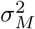 by first inputting into the model 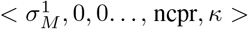 and then sampling from a normal distribution parameterized by the predicted mean and standard deviation of 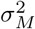. To generate 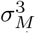, we inputted into the model 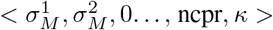 and then sampled from a normal distribution parameterized by the predicted mean and standard deviation of 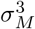. This process is iteratively repeated until the complete output vector < 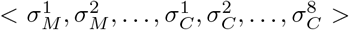 > is generated. For comparison, we also generated samples from the model when we did not condition on ncpr and *κ*. In this case, we sampled a random value of 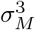 from the training set, and then sampled the remaining singular values according to the same procedure previously outlined.

### 2.6 Reconstructing M and C matrices from samples generated by the model

Because we are only learning the first 8 diagonal entries of Σ_*S, M*_ and Σ_*S, C*_, we used the remaining diagonal entries of Σ_*GS, M*_ and Σ_*GS, C*_ to reconstruct **M** and **C**. Specifically, we combined Σ_*S, M/C*_ with Σ_*GS, M/C*_ by replacing the first eight diagonal entries of Σ_*GS,M/C*_ with that of Σ_*S,M/C*_. We refer to this combined matrix as Σ_*S*+*GS, M/C*_. Generating the reconstructed **M/C** is then achieved via 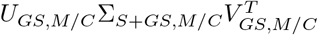.

For reference, we refer to the reconstructed **M/C** as **M**\*/**C**\*. For a given **M**\* and **C**\*, one can also generate samples from a multivariate normal distribution whose mean vector corresponds to **M**\*and whose covariance matrix corresponds to **C**\*. Sampling from this distribution yields a vector of pairwise distances for all pairs of residues. A summary of the overall data pipeline is provided in Fig. 2.

**Figure 2:**
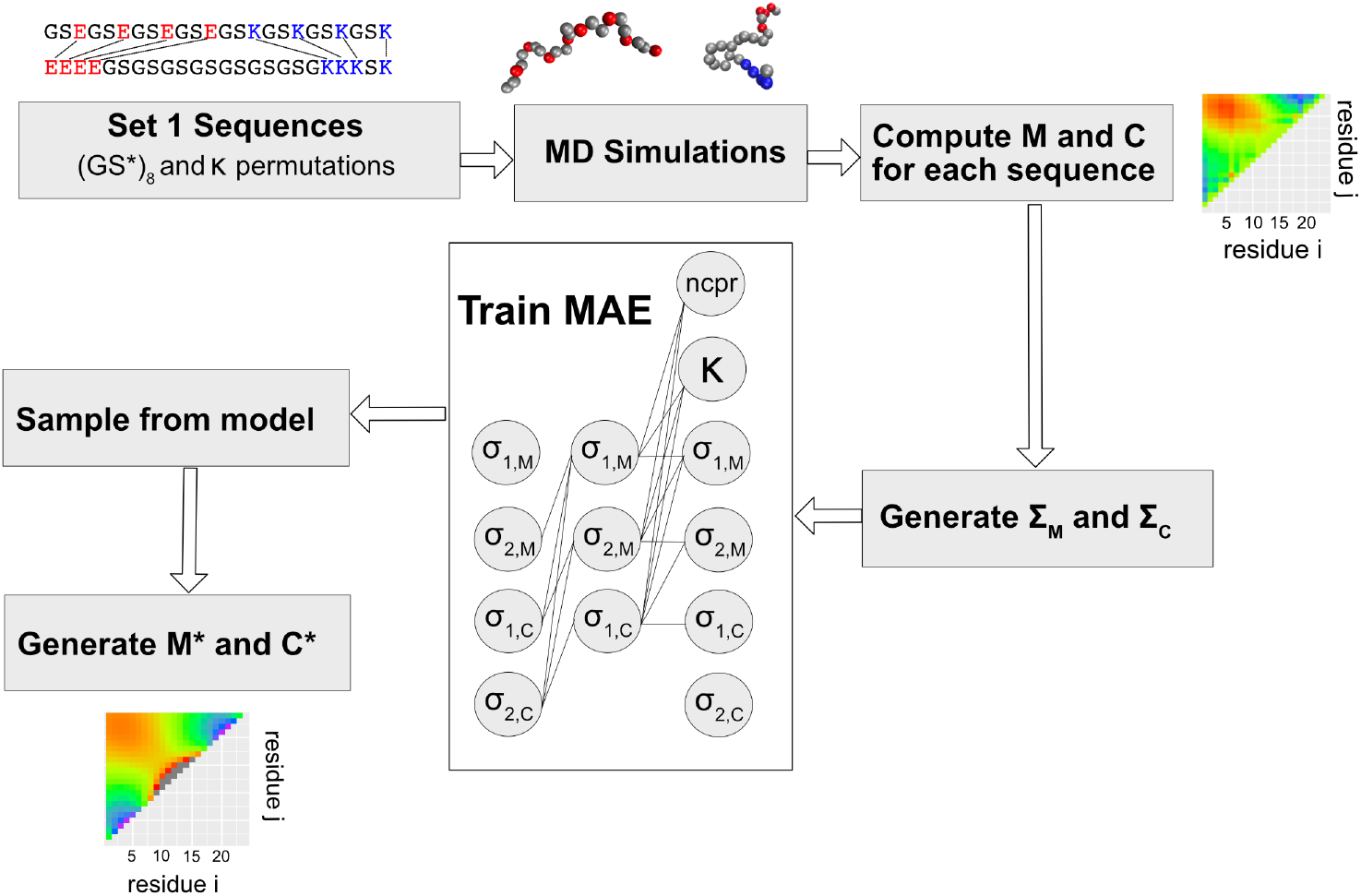
Summary of the overall data pipeline. Coarse-grained molecular dynamics simulations are run on the relevant input sequences. For each sequence, a mean pairwise distance matrix (**M**) and a matrix representing the covariance between each pair of pairwise distances (**C**) is generated. Then, an approximation of the singular values of **M/C** with respect to the (GS)_12_ basis is computed. This input is used to train the conditional masked autoencoder. We can then sample from the model for a given NCPR and *κ* value, and then reconstruct **M** and **C** using the (GS)_12_ basis.

## 3 Results

### 3.1 Evaluating the accuracy of the model

We calculated the test loss of the model with and without conditioning on sequence features (i.e ncpr, *κ*) to quantify their predictive power. In addition, we calculated the test loss of the model with various partitions of the train and test set. For one of the train and test set partitions, we used a standard 80/20 split. Because the training dataset contains 100 instances per sequence (due to the bootstrapping scheme to generate Σ_*GS,M*_ and Σ_*GS, C*_), we created this partition such that a given sequence exists in either the train or test set. The remaining partitions were constructed based on the *κ* value, due to its strong correlation to *R_g_* (spearman correlation coefficient of −.46, Fig 1B). Our rationale was that if a training set contains sequences with a *κ* value in a particular range, the remaining sequences in the test set are more likely to contain ‘unseen’ sequences. We explored various partitions based on *κ*, where sequences in the training set had a *κ* value ranging between [*a, b*] where the difference between *b* and *a* was either .2 or .1. In Fig. 3, we observe that conditioning on sequence features consistently reduced the test loss, and that the model experiences a markedly higher test loss when evaluating sequences whose *κ* value ranged between [0, .1], [.1, .2], and [0, .2] compared to those outside this range. We speculate that because the inverse nonlinear relationship between *κ* and *R_g_* most prominently manifests at lower *κ* values (Fig. 1B), it is more difficult for the model to extrapolate to *κ* values ranging between [0, .2] as opposed to [.6, .8]. Finally, we note that the model incurs the lowest test loss for *κ* values ranging between [.3, .6], suggesting the model has a strong capability to interpolate.

**Figure 3:**
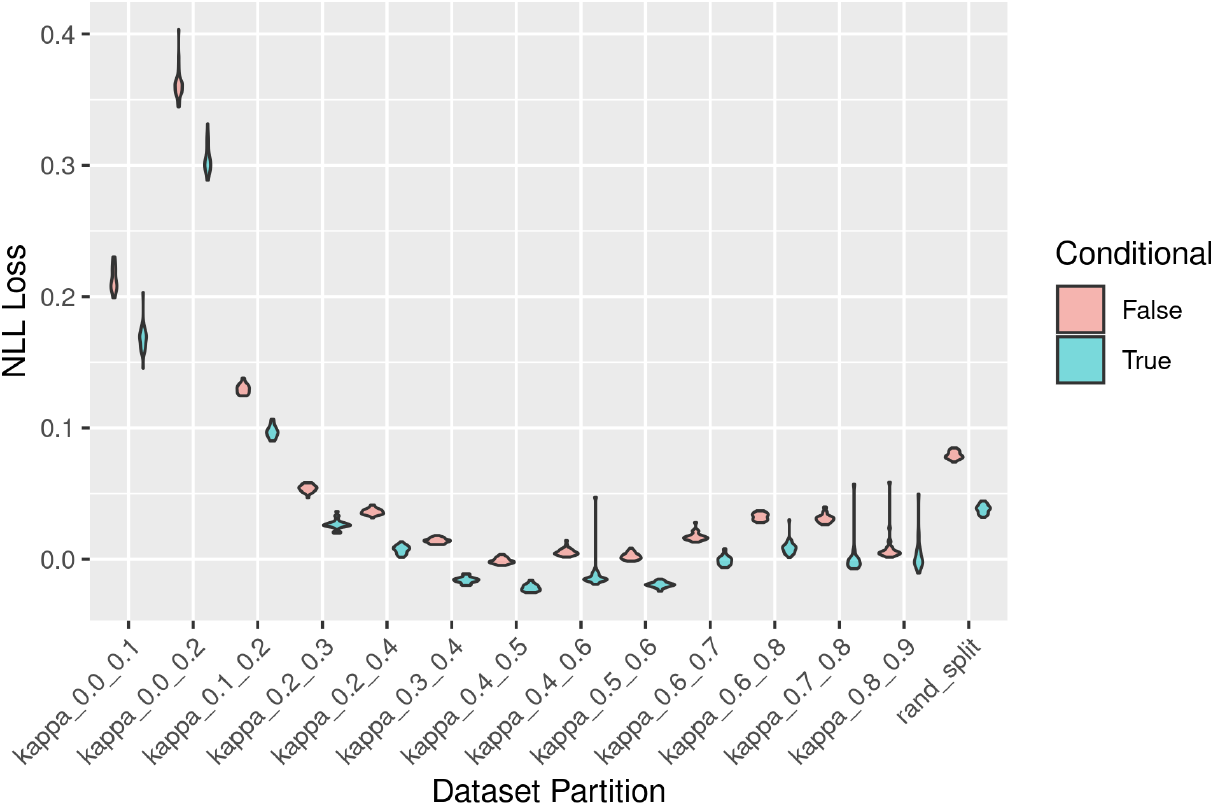
Negative likelihood loss distribution for different train/test partitions. The x-axis refers to a particular train/test set partition. X-axis labels of the form “kappa_a_b” mean that the train set consisted of all sequences whose *κ* value was outside the window [*a, b*], while the test set consisted of all sequences whose *κ* value was within [*a, b*]. The x-axis label “rand_split” refers to the 80/20 split, where a given sequence was in either the train or test set. For each train/test partition, we repeated the NLL loss calculation 30 times, as opposed to providing solely a point estimate. Finally, we evaluated the effect of training the model with (blue) and without (red) conditioning on sequence features.

### 3.2 Evaluating the singular value distribution generated by the model

In Fig. S13-15, we visualize the two-dimensional distribution of a subset of variables in < *x, c* > for samples generated by the model and samples from set 1. We specifically overlay the two dimensional distribution of samples generated by a given model whose test set consisted of sequences whose value ranged between [*a, b*] and samples from from set 1 whose *κ* value ranged between [*a, b*]. At a qualitative level, we observe that the model has a strong ability to recapitulate pairwise relationships between variables in < *x, c* >. To quantify this trend more precisely, we calculated the absolute value of the difference in the spearman correlation between each pair of variables in < *x, c* > for samples generated by the model where *κ* was restricted to values in the test set and the appropriate samples in set 1 (Fig. 4). For context, we also compared 1) samples generated by the same model when no restrictions were placed on the value of *κ* to samples from the entire dataset and 2) samples generated by the same model when the model did not condition on sequence features to samples from the entire dataset.

**Figure 4:**
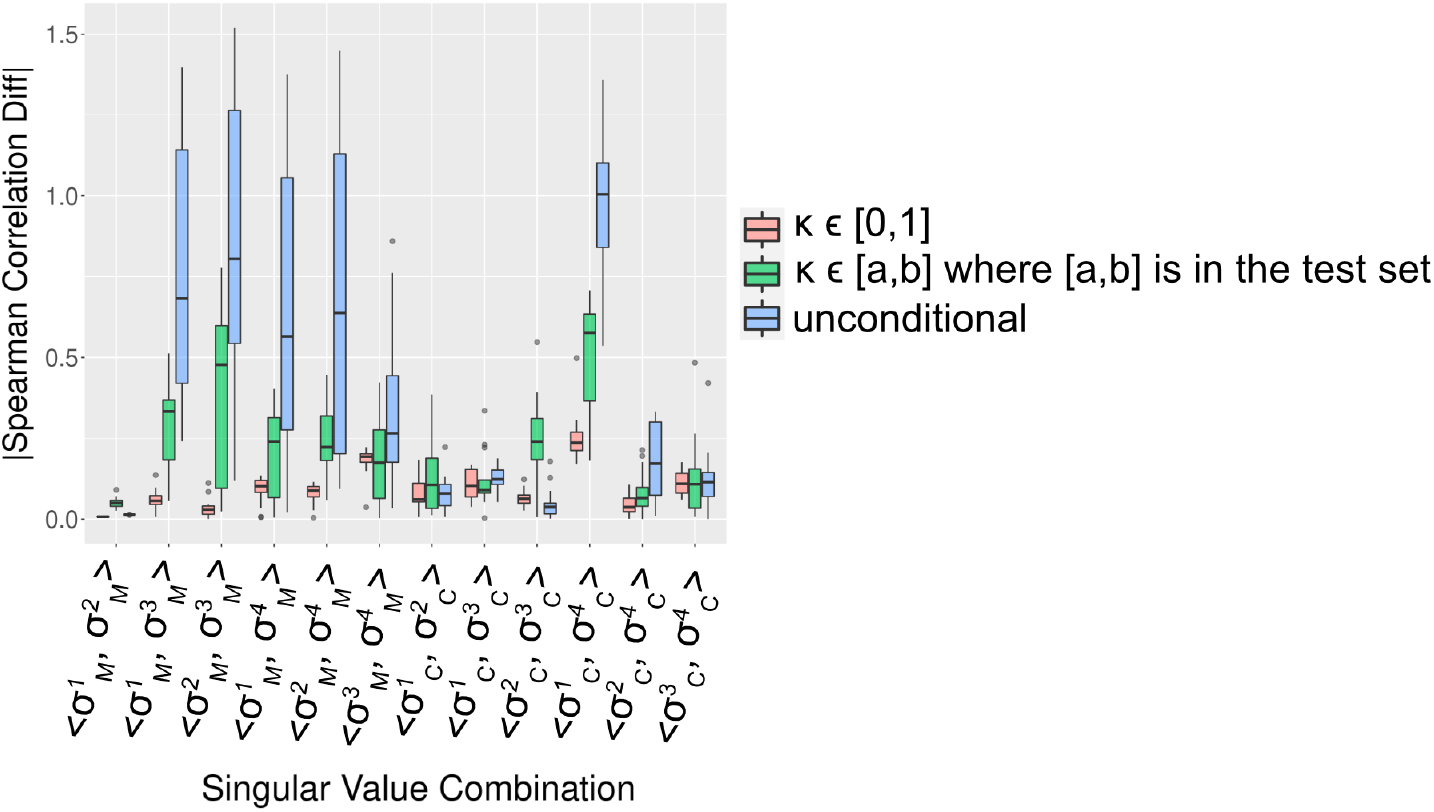
Quantifying the difference between the distribution of pairs of variables in < 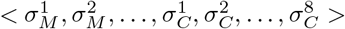 > for samples in set 1 and samples generated by the model. The x-axis for each panel refers to the specific combination of pairwise variables. The y-axis refers to the absolute value in the difference between the spearman correlation. For each singular value combination, we report the error across all *κ*-based train/test partitions. Specifically, for each model derived from a specific *κ*-based train/test partition, we calculated the error metrics between samples generated by the model where *κ* was restricted to values in the test set and samples from set 1 whose *κ* value is in the test set (green). We also made two sets of additional comparison: 1) samples generated by the model when no restrictions were placed on the value of *κ* versus all samples in set 1 (red), and 2) samples generated by the model when the model did not condition on sequence features versus all samples in set 1 (blue).

We observe that samples generated by the model when no restrictions were placed on the value of *κ* consistently have the lowest errors, suggesting the models ability to fit the training data well. Furthermore, the error for samples generated by the model where *κ* was restricted to values in the test set are relatively low and comparable to that of samples generated without any *κ* restrictions. Finally, we note that samples generated by the model when the model did not condition on sequence features consistently have the highest errors, indicating the predictive power of sequence information when modeling the distribution of 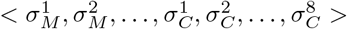.

### 3.3 Average pairwise distance maps and two-dimensional pairwise distance distributions generated by the model

Given a set of singular values generated by the model, we generated their corresponding **M**\* and **C**\* matrices according to the procedure outlined in Methods. We first compare sets of **M**\* generated by different models trained under different train/test partitions to sets of **M** from the test set partition (Fig. 5, Fig. S16). For visual clarity, we normalized each **M** and **M**\* by dividing it by **M**_*GS*_. Overall, the generated sets of **M**\* qualitatively capture the trends of **M** from the test set partition. However, they do not appear to completely capture the heterogeneity and richer variation present across sets of **M** from the test set partition. For a given **M**\* and **C**\*, we also generated 300 samples from 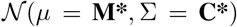 and plotted two-dimensional pairwise distance distributions for a subset of residue pairs (i.e < *d*ij**, *d_kl_* >, where < *i, j* >and < *k, l* > correspond to arbitrary pairs of residues). In Fig. S17, we provide a reference point and visualize the two-dimensional pairwise distance distributions derived from trajectory data for a sequence with a *κ* value of .3. In Fig. S18, we visualize the two-dimensional pairwise distance distributions generated by a model whose test set consisted of sequences whose *κ* value ranged between [.2, .4]. Similar to the observation noted for the generated sets of **M**\*, we observe a generally strong agreement between two-dimensional pairwise distance distributions that were generated via the model and from actual sequences in the testing set.

**Figure 5:**
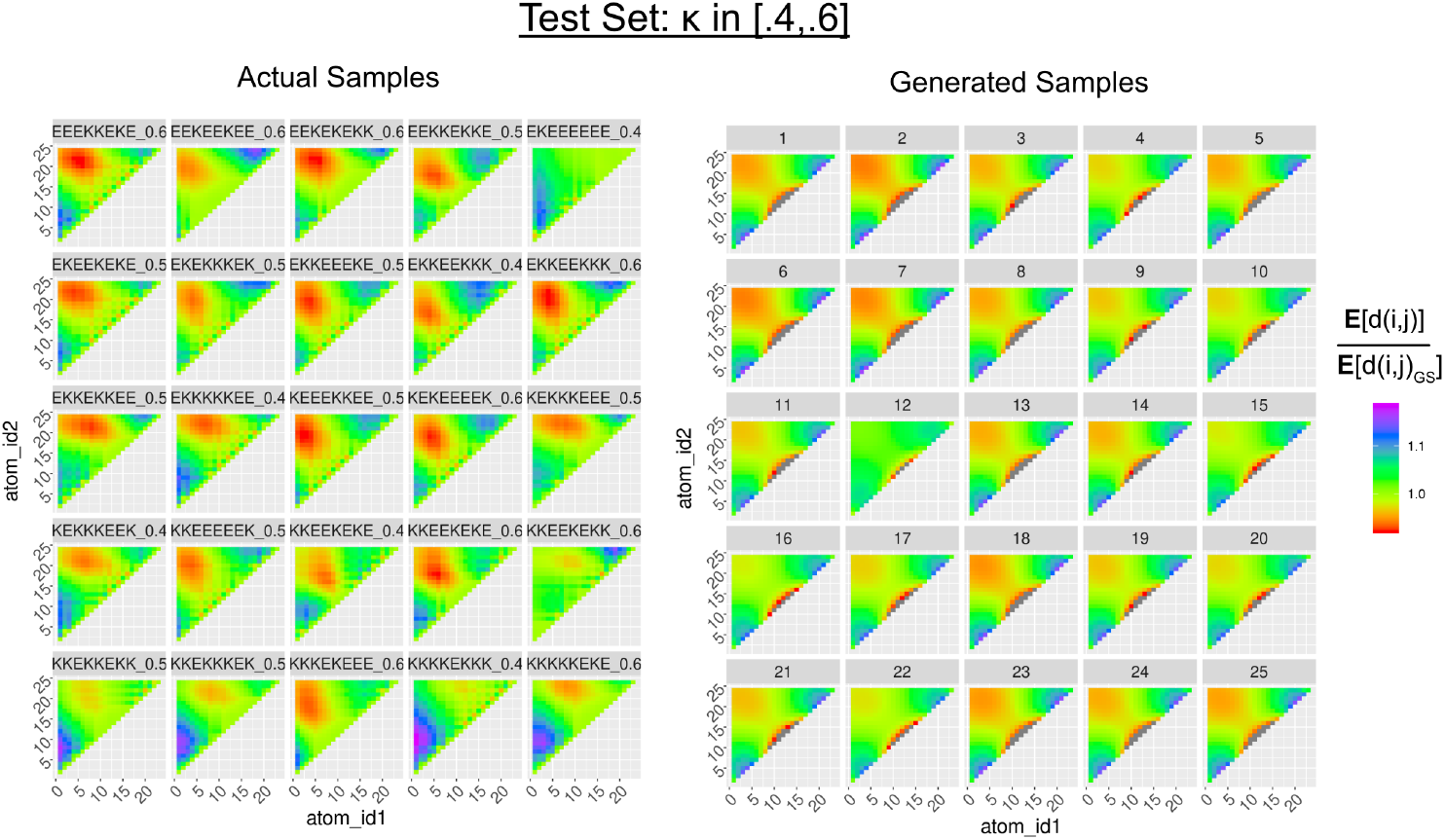
On the left, are actual samples from the *κ* permutations set while on the right are samples generated by the model. The model was trained on inputs whose *κ* values were outside the range [.4, .6]. Note that to sample from the model, *κ* was restricted between [.4, .6]. In the distance map, each value represents the average distance between residue *i* and *j* divided by the average distance between residue i and j from the (GS)_12_ simulation. Finally, we note that the color scale is based on the minimum and maximum value among the panel of plots of actual samples. To maintain ease of comparison, any value derived from the generated samples below or above the corresponding minimum or maximum value was set to gray.

### 3.4 Comparing the singular value distribution of sequences in the training set to sequences with heterogeneous compositions

Naturally, one may ask whether this approach generalizes to sequences of arbitrary composition. To test this, we compared the pairwise distribution of singular values for sequences consisting only of glycine, serine, glutamate, and lysine to the *κ* variants and hydrophobic variants described in the second subsection of methods (Fig. S19 and S20). Though there is some overlap between the pairwise distributions (most prominently reflected in 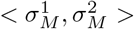), we overall observe a significant degree of deviation. We note this deviation is not due to the use of the (GS)_12_ basis as the reconstruction error of **M** is comparable to that of sequences in set 1 (Fig. S21 and S22). Though this trend may be unsurprising for the hydrophobic variants, the fact that it also holds for the *κ* variants suggests that heterogeneity in sequence composition leads to unique fingerprints of pairwise distance distributions despite homogeneity in sequence features. From a biophysical perspective, it may be unremarkable to note that a set of sequences with heterogeneous composition but homogeneous sequence features can have different biophysical behavior. However, quantifying that difference in a low-dimensional space (i.e 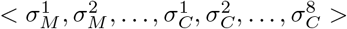) provides a relevant, complementary context.

## 4 Conclusion

In this work, we demonstrated an approach to train a masked auto-encoder to learn the parameters of a multivariate normal distribution modeling pairwise distance information derived from simulations of short, disordered proteins. This was achieved by representing the spatial output of a molecular dynamics simulation in a low-dimensional space, and training the model to learn the distribution of this encoding. Furthermore, by sampling from the masked auto-encoder and applying the relevant inverse transformations, one can generate realistic, averaged pairwise distance maps. This suggests there are learnable patterns in structural features of disordered proteins. Though the data originating from the *κ* and hydrophobic variants were partially out-of-distribution, we speculate that our approach can be applied to disordered sequences of longer length and more heterogeneous composition, given the appropriate training set. Relevant data and code is available at https://www.dropbox.com/sh/4zz1i4zgoaxhjf0/AAATNiuEAn9rFJ-Tqhzjr-zpa?dl=0 and https://github.com/itaneja2/idr_design_w_data respectively.

