## Supplementary Information for "Generative modeling of short, disordered proteins with homogeneous sequence composition"

### Supplementary Figures

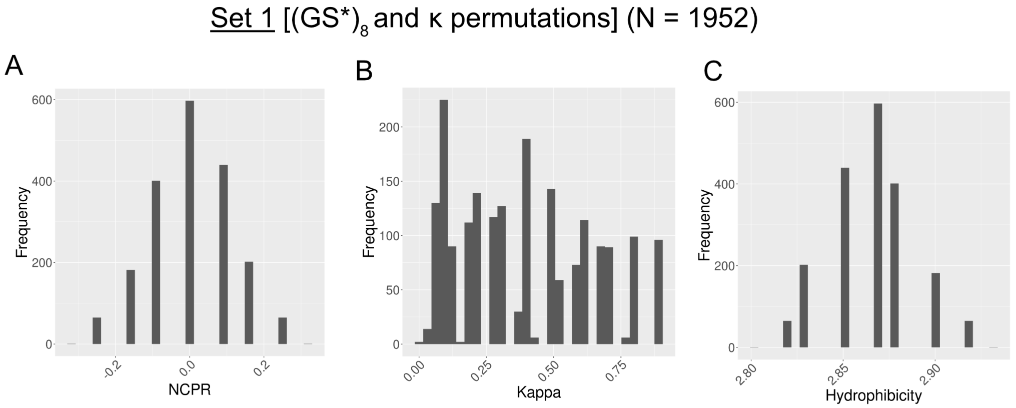


Figure S1. **Sequence properties in the (GS*)$_8$ and $\kappa$ permutations set.** Histograms for the net charge-per residue (A), $\kappa$ (B), and hydrophobicity (C). For reference, the hydrophobicity is defined as the mean hydropathy of a sequence, calculated as the average of a 0-9 renormalized Kyte-Doolittle hydrophobicity scale.

Set 2 ($\kappa$ variants)

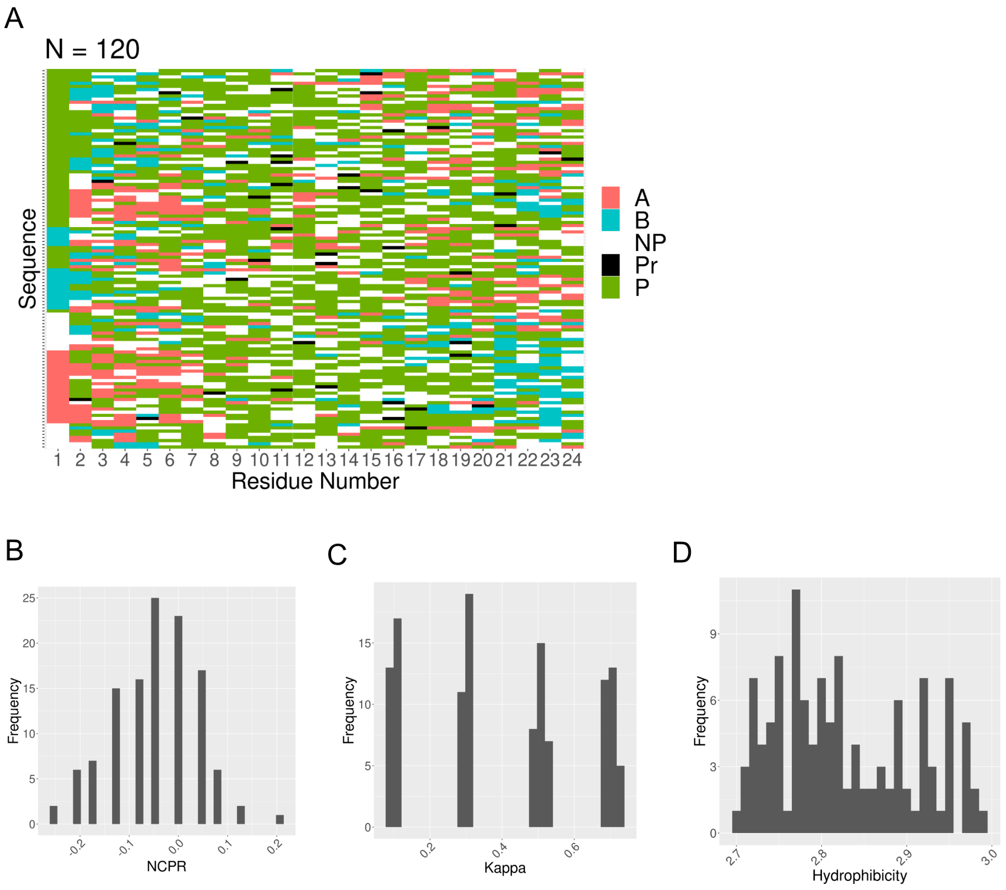


Figure S2. **Sequence properties in the $\kappa$ variants set, where $\kappa$ was restricted to values in the set $\{.1, .3, .5, .7\}$.** For (A), the y-axis corresponds to a given sequence in the $\kappa$ variants set. For each sequence, its residue is color coded according to its category (Acidic = Red, Basic = Blue, Non-polar = White, Proline = Black, and Polar = Green). We also plot histograms for the net charge-per residue (B), $\kappa$ (C), and hydrophobicity (D). For reference, the hydrophobicity is defined as the mean hydropathy of a sequence, calculated as the average of a 0-9 renormalized Kyte-Doolittle hydrophobicity scale.

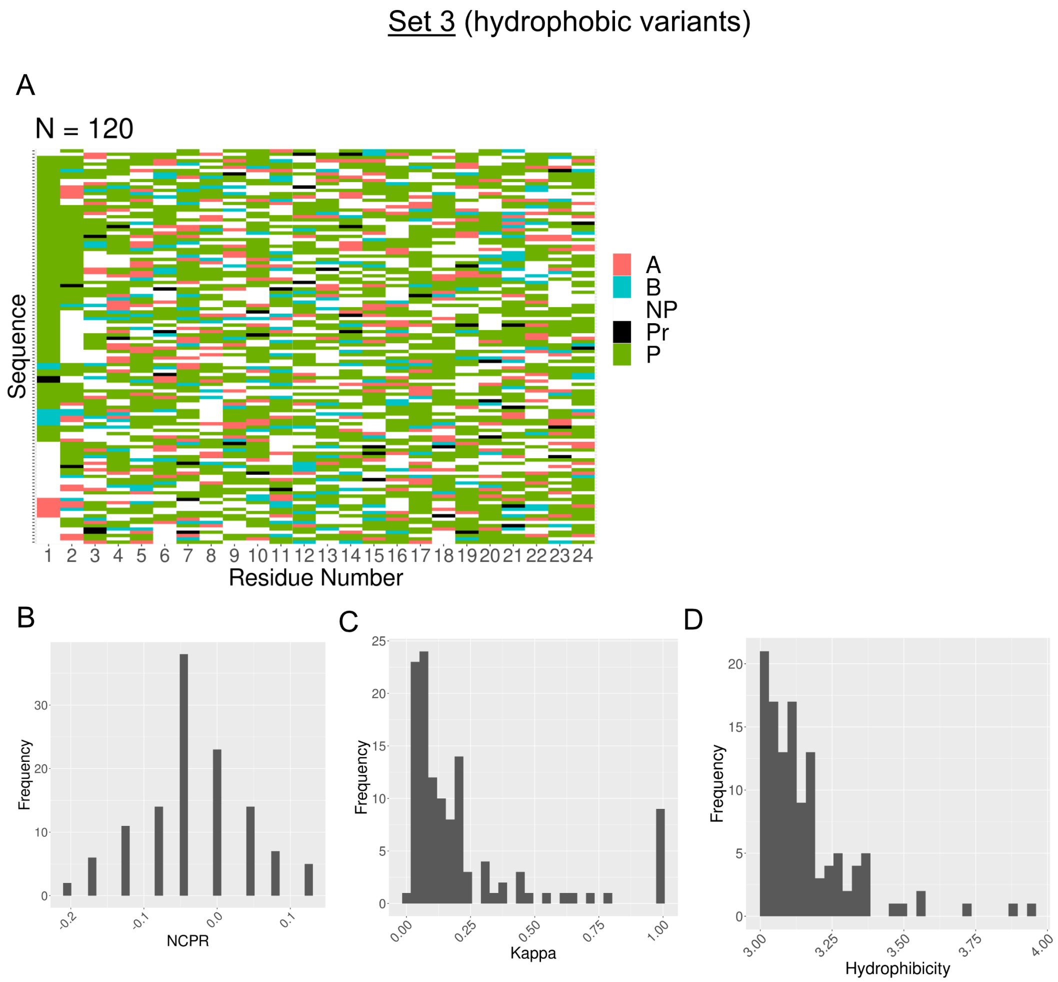


Figure S3. **Sequence properties in the hydrophobic variants set.** For (A), the y-axis corresponds to a given sequence in the hydrophobic variants set. For each sequence, its residue is color coded according to its category (Acidic = Red, Basic = Blue, Non-polar = White, Proline = Black, and Polar = Green). We also plot histograms for the net charge-per residue (B), $\kappa$ (C), and hydrophobicity (D). For reference, the hydrophobicity is defined as the mean hydropathy of a sequence, calculated as the average of a 0-9 renormalized Kyte-Doolittle hydrophobicity scale.

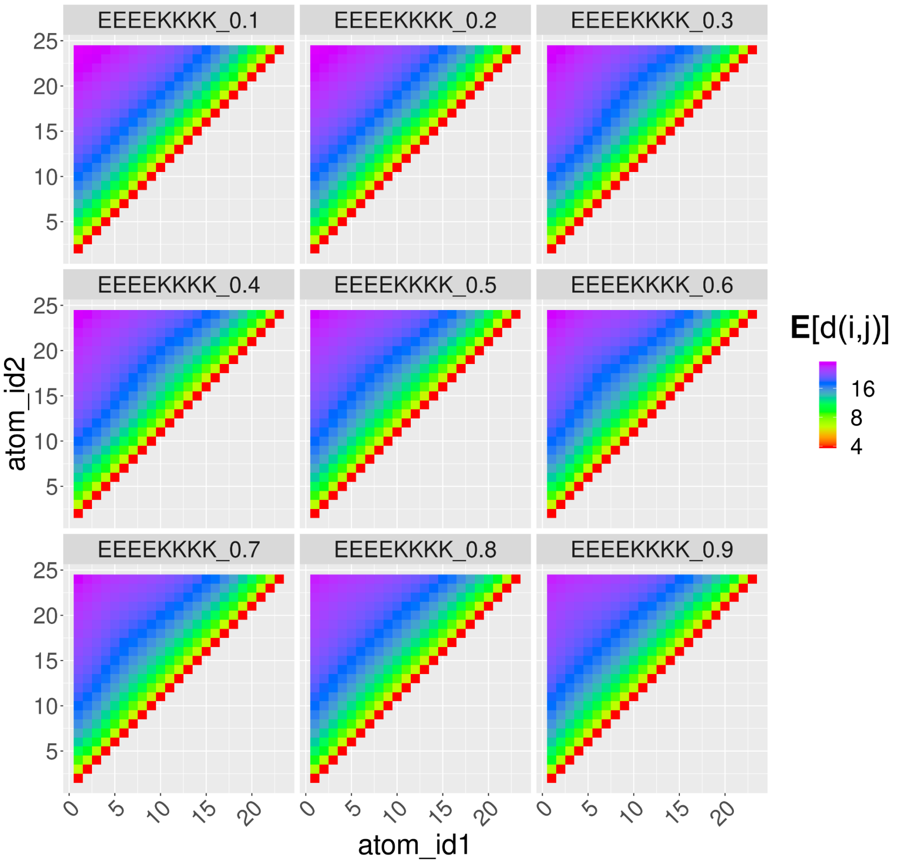


Figure S4. **Ensemble-averaged pairwise distance maps for a subset of sequences in the $\kappa$ permutations set.** Each value in the distance map represents the average distance between residue $i$ and $j$.

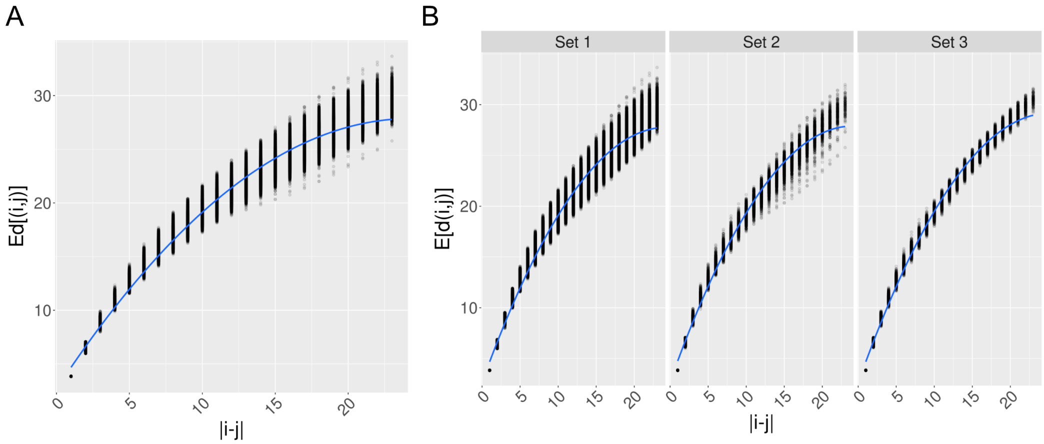


Figure S5. **Correlation between $|i - j|$ and $\mathbf{E}[d(i, j)]$.** A) Correlation between $|i - j|$ and $\mathbf{E}[d(i, j)]$ among all sets of sequences (1, 2, and 3). B) Correlation between $|i - j|$ and $\mathbf{E}[d(i, j)]$ for each set of sequences.

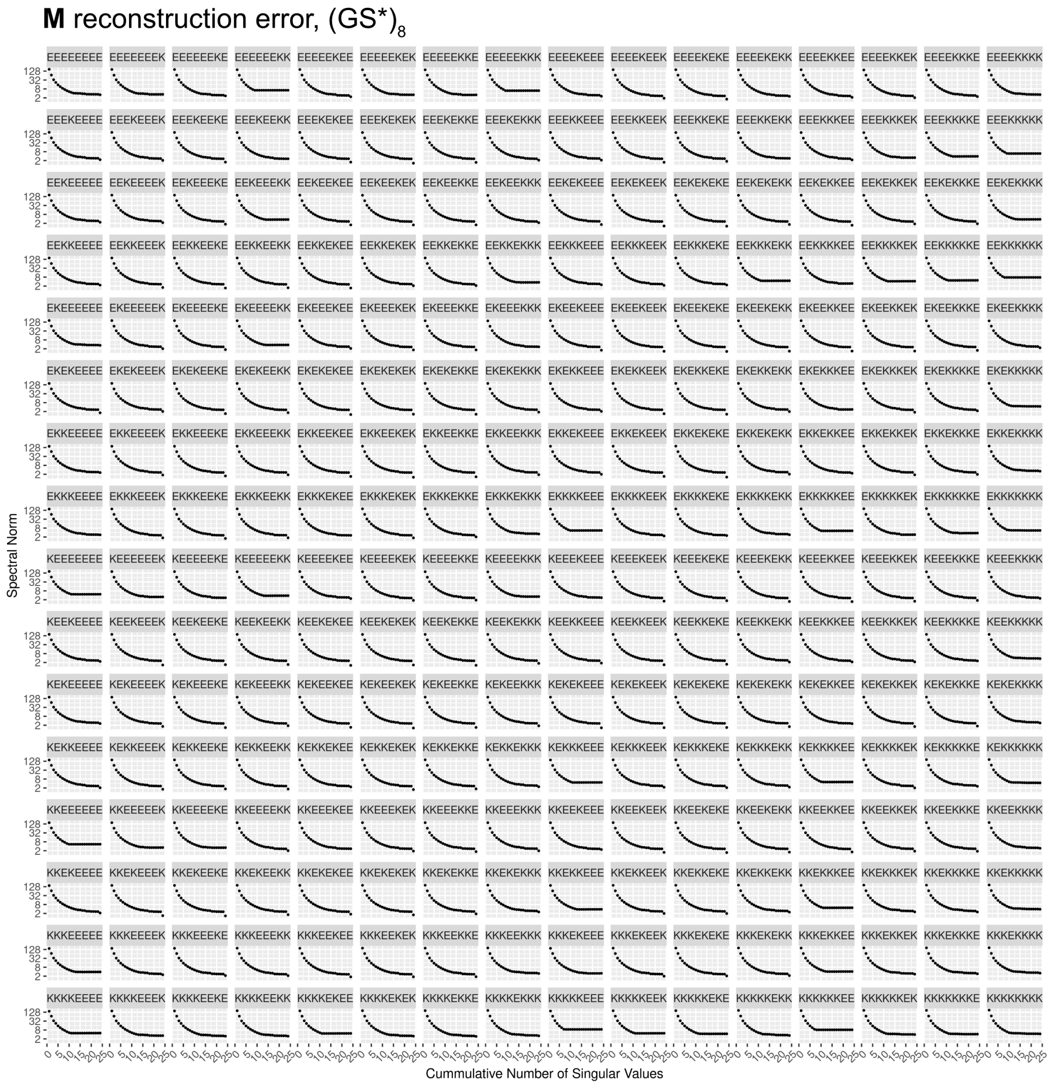


Figure S6. **Reconstruction error of M for each sequence in the $(GS^*)_8$ set as a function of the number of cumulative singular values used.** Reconstruction error is defined as the spectral norm between the difference of the reconstructed matrix and original matrix.

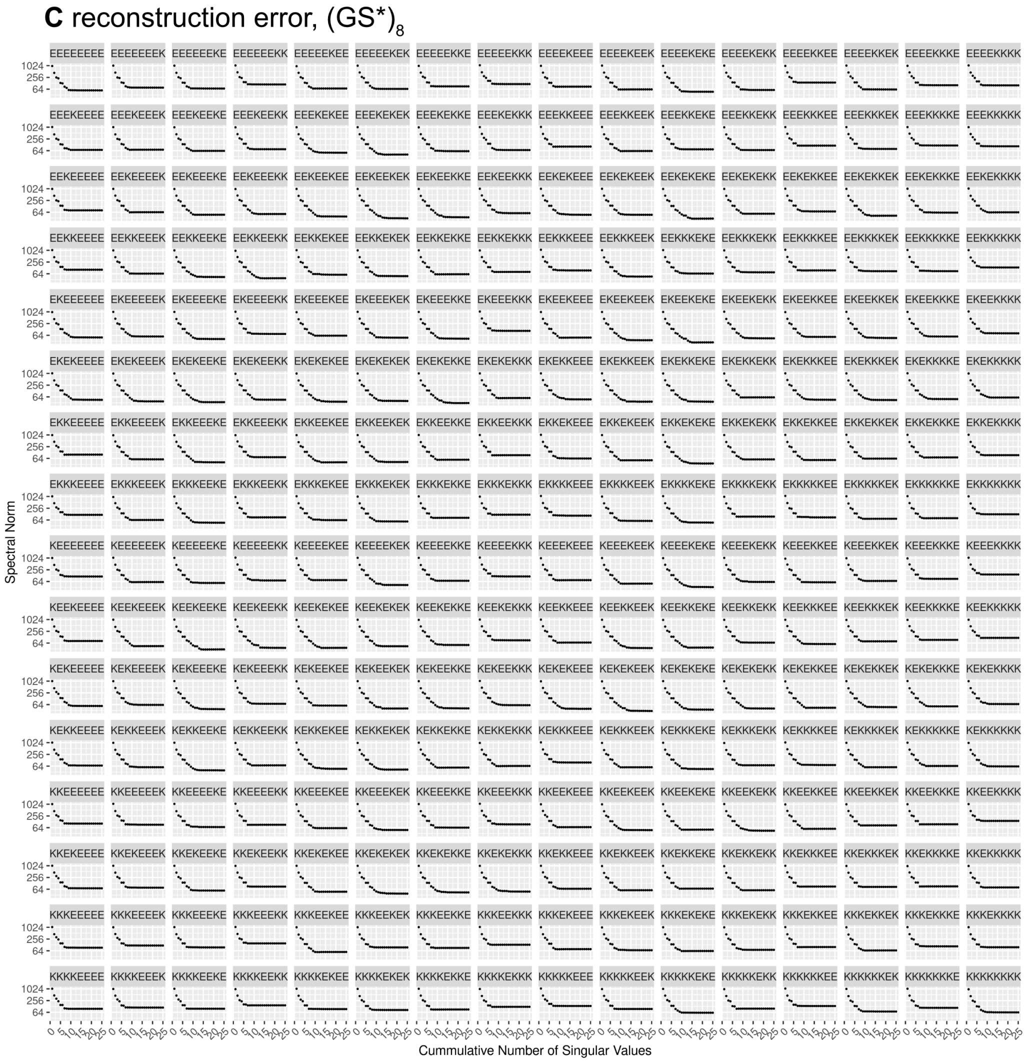


Figure S7. **Reconstruction error of C for each sequence in the $(GS^*)_8$ set as a function of the number of cumulative singular values used.** Reconstruction error is defined as the spectral norm between the difference of the reconstructed matrix and original matrix.

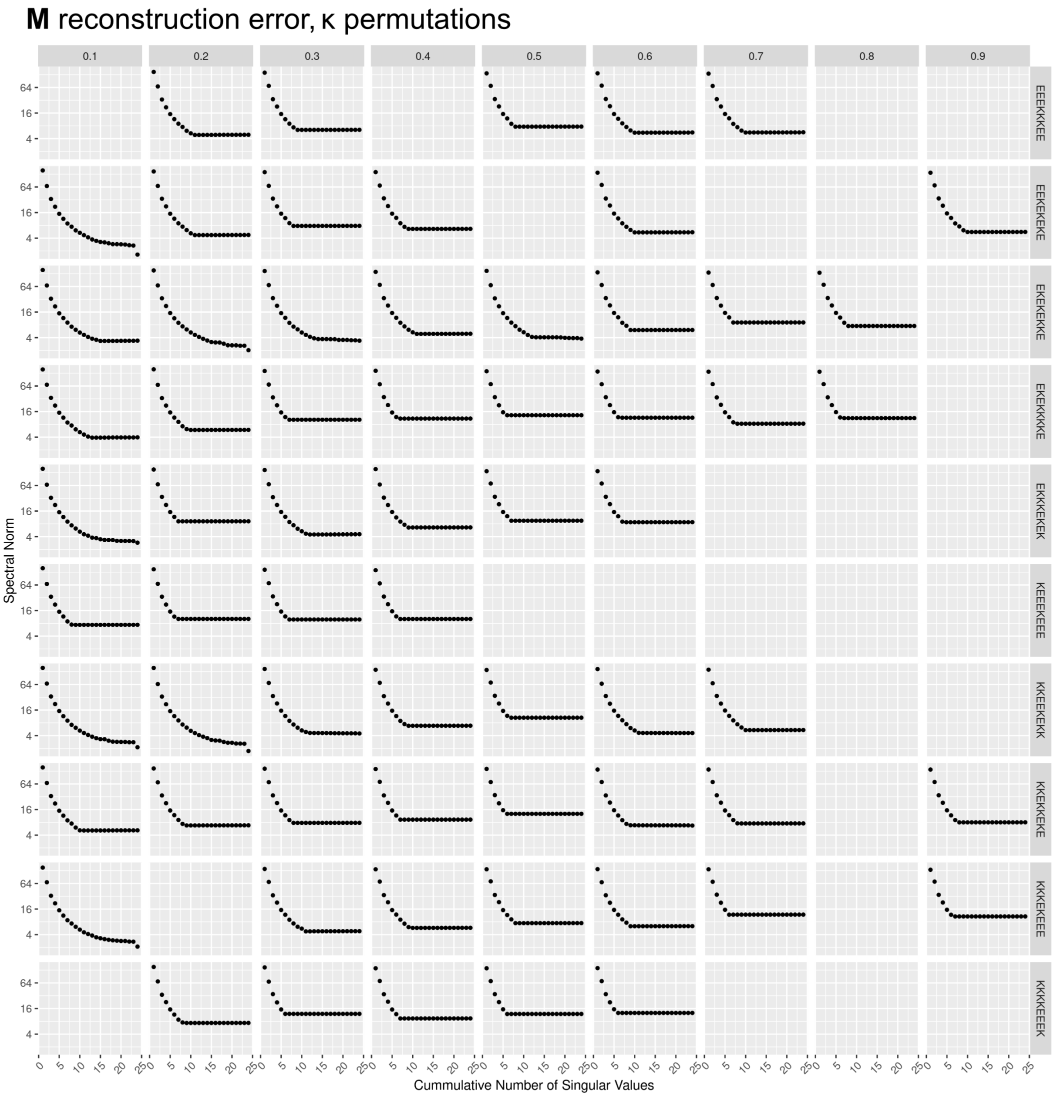


Figure S8. **Reconstruction error of M for a subset of sequences in the $\kappa$ permutations set as a function of the number of cumulative singular values used.** Reconstruction error is defined as the spectral norm between the difference of the reconstructed matrix and original matrix. Each column refers to a certain value of $\kappa$ while each row corresponds to the sequence from which the permuted variant was derived from.

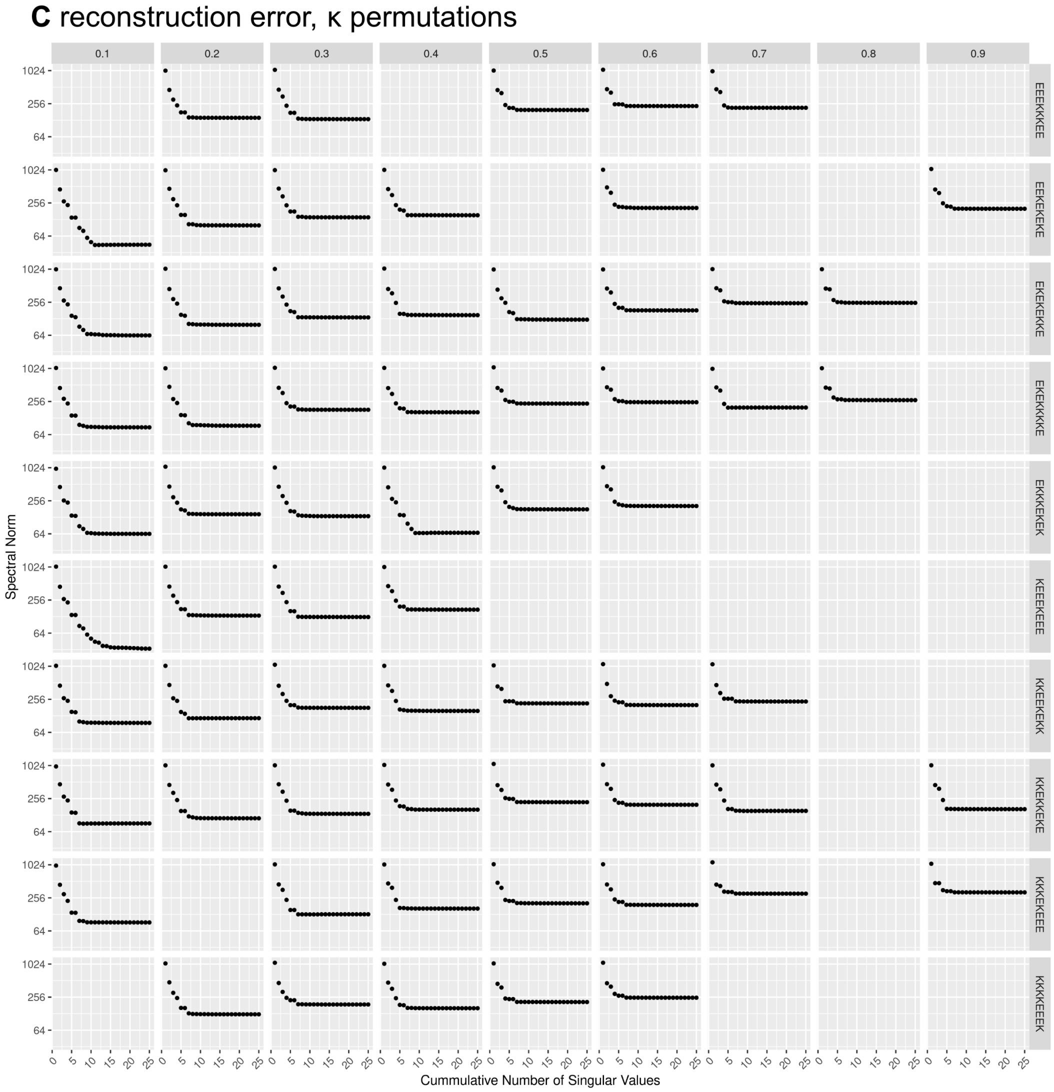


Figure S9. **Reconstruction error of C for a subset of sequences in the $\kappa$ permutations set as a function of the number of cumulative singular values used.** Reconstruction error is defined as the spectral norm between the difference of the reconstructed matrix and original matrix. Each column refers to a certain value of $\kappa$ while each row corresponds to the sequence from which the permuted variant was derived from.

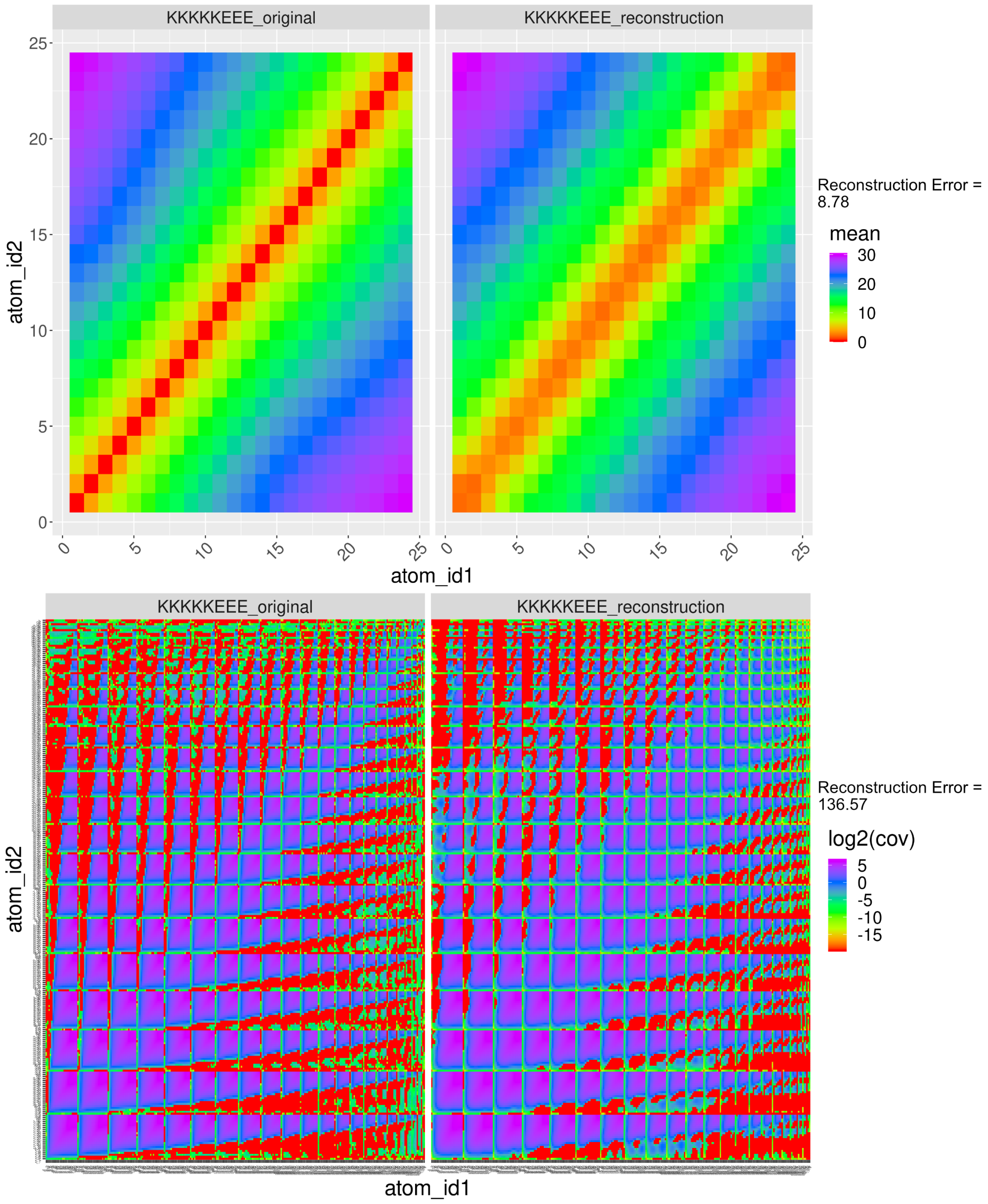


Figure S10. **Comparison of the reconstructed M (top-right) or C (bottom-right) and original M (top-left) or C (bottom-left) for a particular sequence in the (GS*)$_8$ set.** For reference, the reconstructed **M** and **C** were derived from $< \sigma_M^1, \sigma_M^2, \ldots, \sigma_C^1, \sigma_C^2, \ldots, \sigma_C^8 >$ respectively. This specific sequence (GSKGSKGSKGSKGSKGSEGSEGSE) was chosen as it has a relatively high reconstruction error for **M** and **C**. We observe a strong correspondence between the reconstructed **M/C** and original **M/C**.

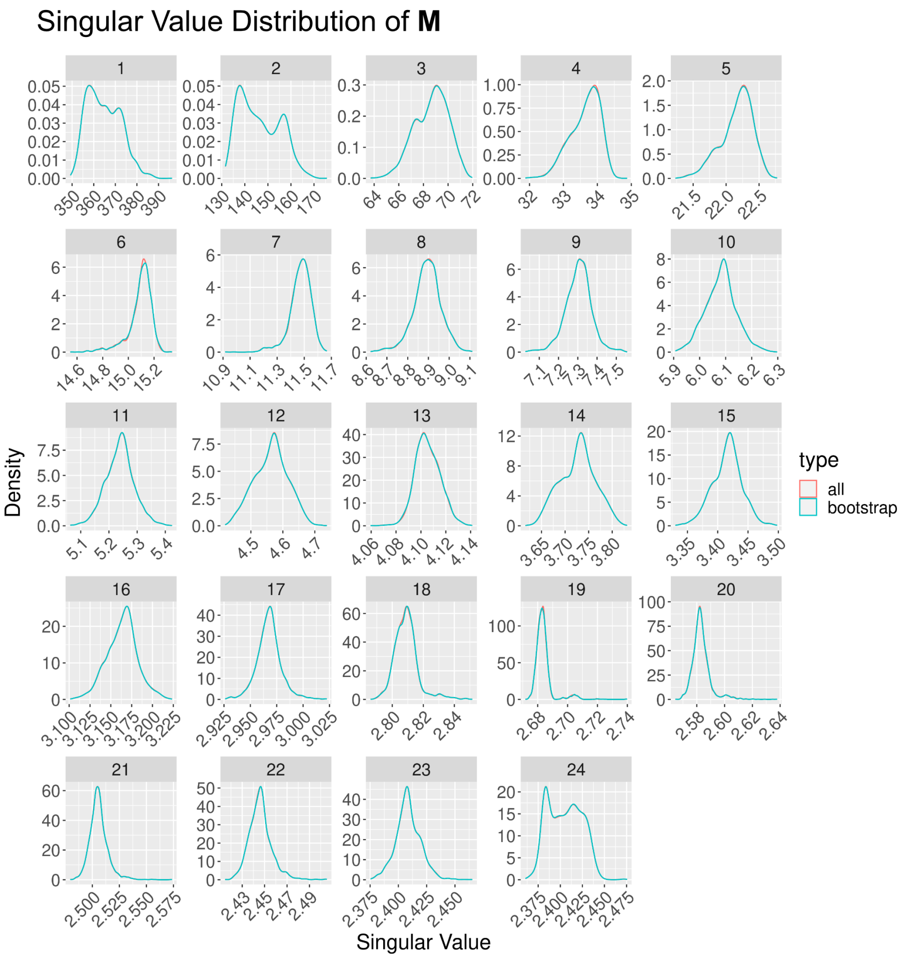


Figure S11. **Distribution of $< \sigma_M^1, \sigma_M^2, \ldots, \sigma_M^{24} >$ when M and C are derived from all frames in the trajectory versus 1000 frames sampled randomly with replacement (i.e bootstrap).** We observe that the distribution derived from all frames in the trajectory and the subsampled frames are nearly equivalent.

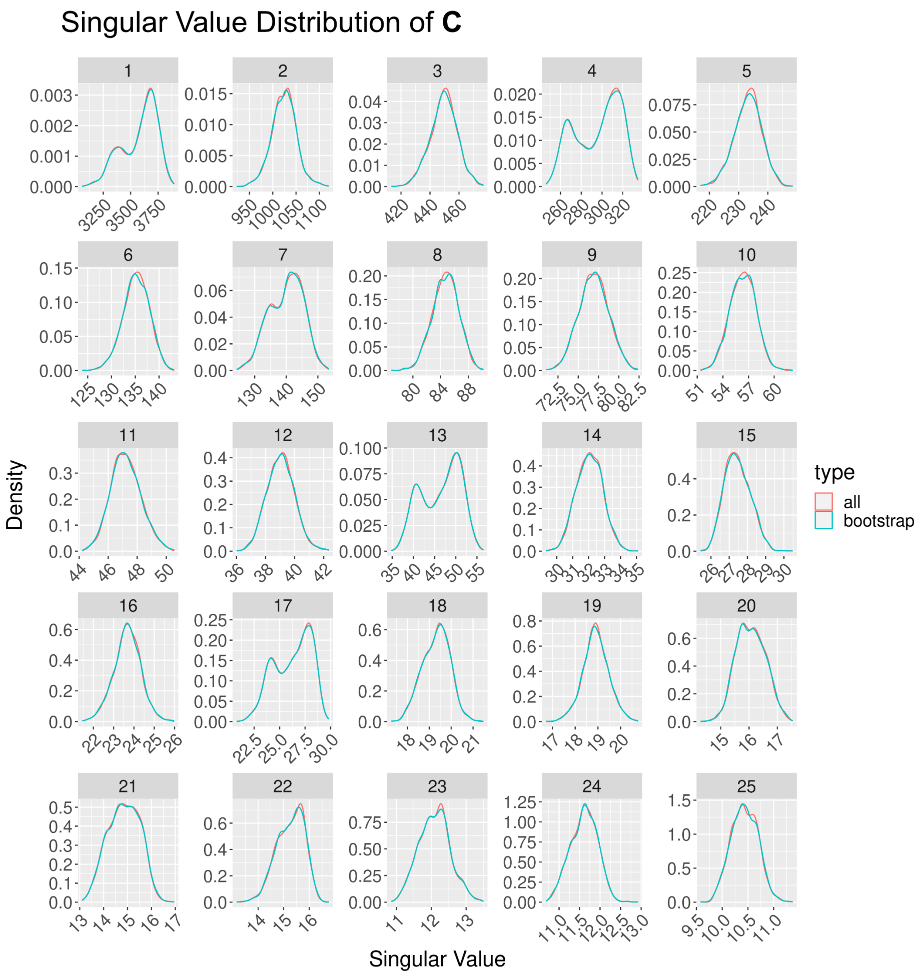


Figure S12. **Distribution of $< \sigma_C^1, \sigma_C^2, \ldots, \sigma_C^{25} >$ when M and C are derived from all frames in the trajectory versus** 1000 **frames sampled randomly with replacement (i.e bootstrap).** We observe that the distribution derived from all frames in the trajectory and the subsampled frames are nearly equivalent. For visual clarity, we only showed the first 25 singular values as opposed to all 276.

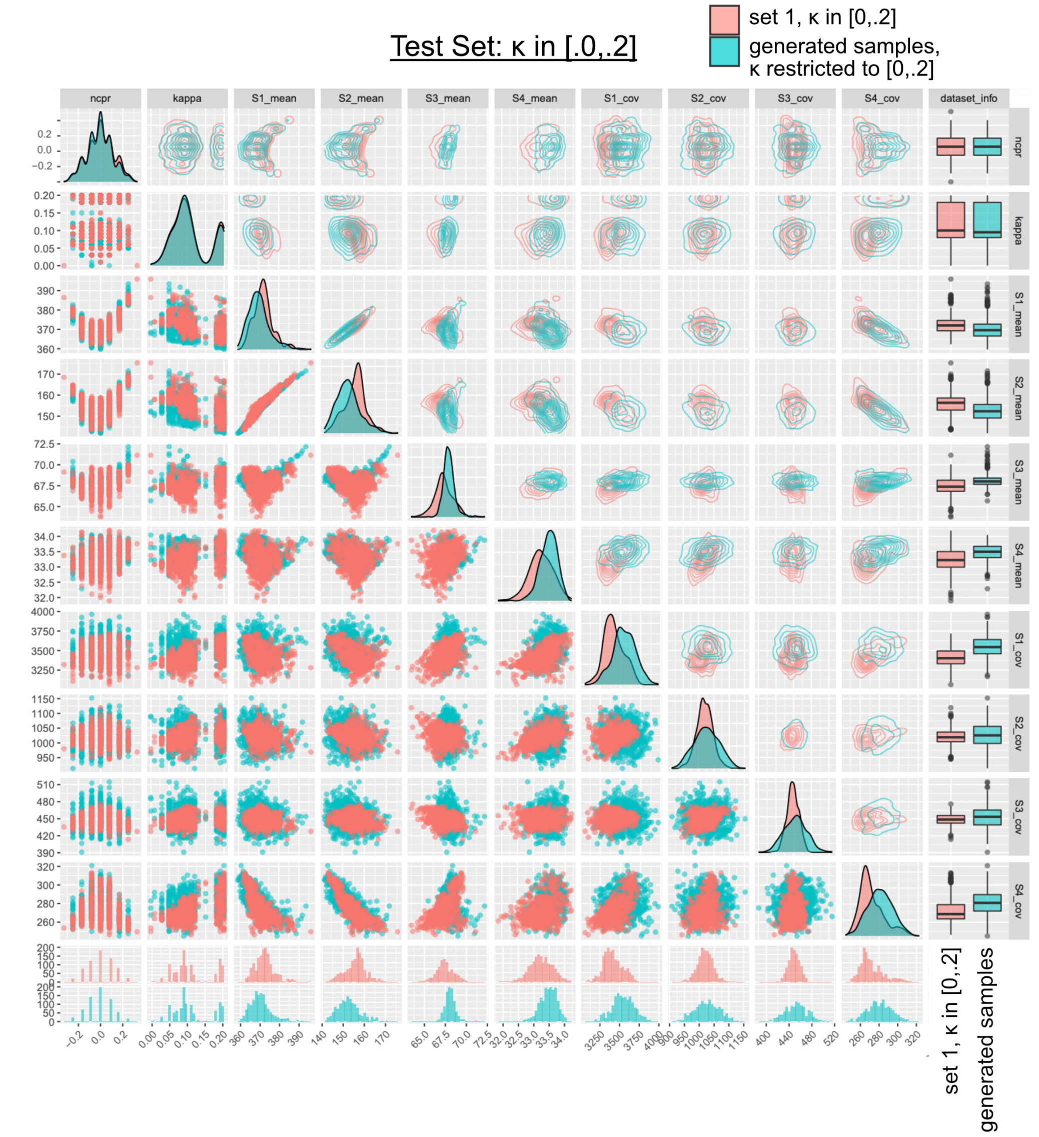


Figure S13. **Pairwise distribution for each pair of variables in $<\sigma_M^1, \sigma_M^2, \sigma_M^3, \sigma_M^4, \sigma_C^1, \sigma_C^2, \sigma_C^3, \sigma_C^4$, ncpr, $\kappa >$ for a subset of samples in set 1 and samples generated by the model.** The model was trained on inputs whose $\kappa$ values were outside the range $[0, .2]$. Points that are colored red refer to data from set 1 whose $\kappa$ value is within $[0, .2]$, and points colored blue refer to data generated by the model. Note that to sample from the model, $\kappa$ was restricted between $[0, .2]$. For reference, the number of instances in the actual and generated sample set is equal. Overall, we observe that the pairwise distributions of the generated samples strongly overlap with that of the actual samples. The x/y-axis of each plot stems corresponding column/row facet. Entries in the lower-triangular portion of the matrix represent scatter plots for each pair of variables while entries in the upper-triangular portion of the matrix represent 2d density contour plots. Entries on the diagonal represent 1d density plots.

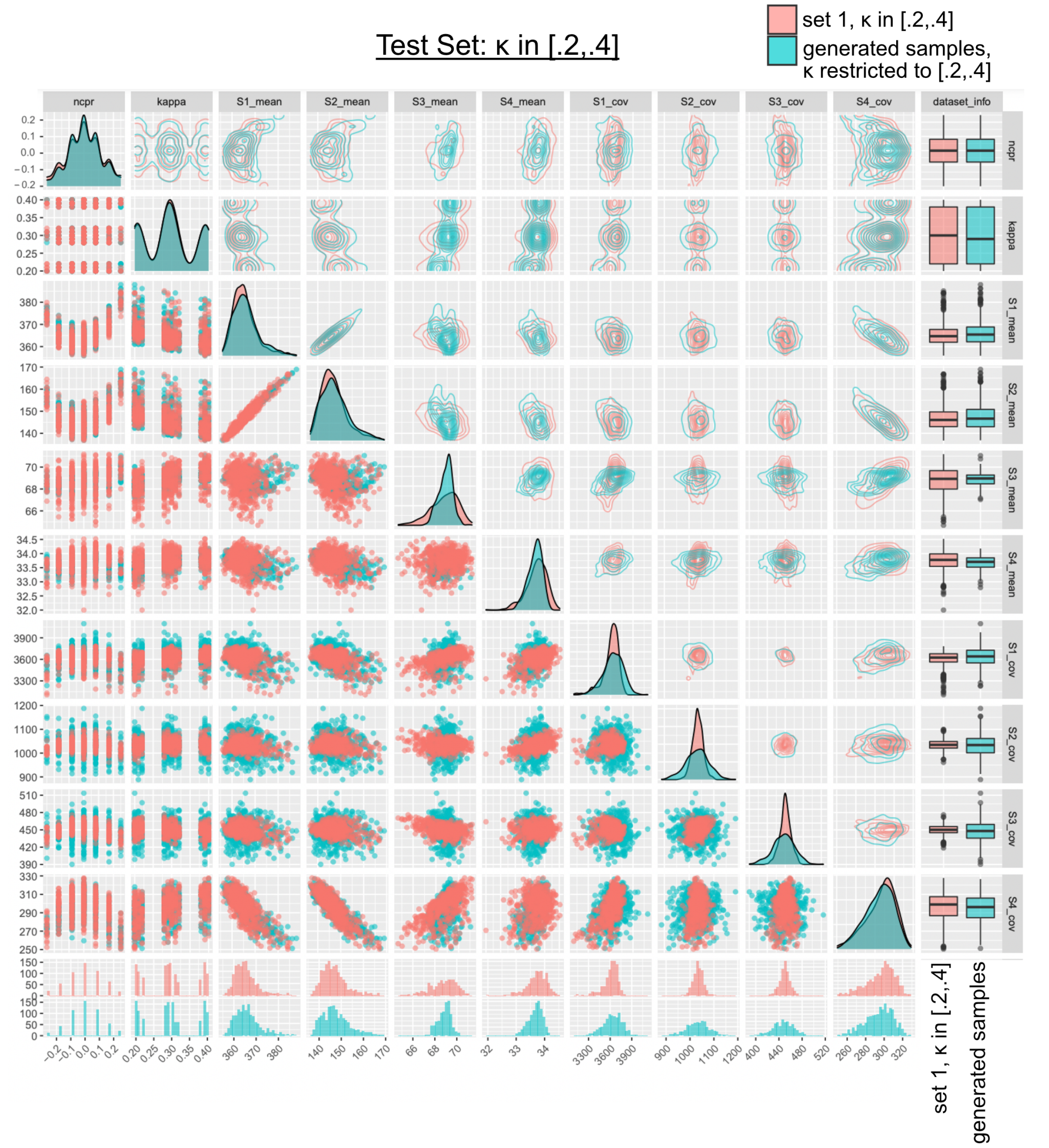


Figure S14. **Pairwise distribution for each pair of variables in $<\sigma_M^1, \sigma_M^2, \sigma_M^3, \sigma_M^4, \sigma_C^1, \sigma_C^2, \sigma_C^3, \sigma_C^4$, ncpr, $\kappa >$ for a subset of samples in set 1 and samples generated by the model.** The model was trained on inputs whose $\kappa$ values were outside the range $[0, .2]$. Points that are colored red refer to data from set 1 whose $\kappa$ value is within $[2, .4]$, and points colored blue refer to data generated by the model. Note that to sample from the model, $\kappa$ was restricted between $[2, .4]$. For reference, the number of instances in the actual and generated sample set is equal. Overall, we observe that the pairwise distributions of the generated samples strongly overlap with that of the actual samples. The x/y-axis of each plot stems corresponding column/row facet. Entries in the lower-triangular portion of the matrix represent scatter plots for each pair of variables while entries in the upper-triangular portion of the matrix represent 2d density contour plots. Entries on the diagonal represent 1d density plots.

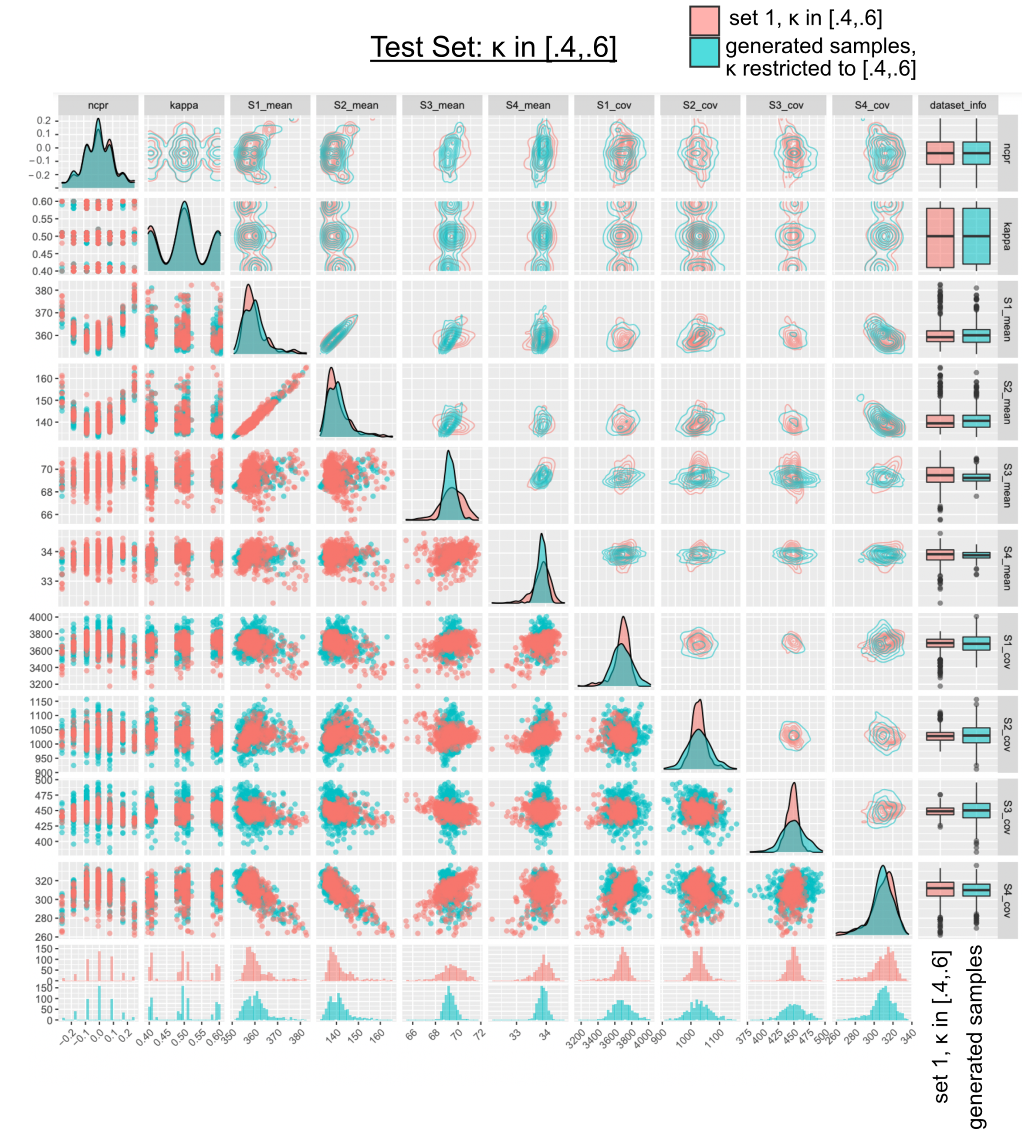


Figure S15. **Pairwise distribution for each pair of variables in $<\sigma_M^1, \sigma_M^2, \sigma_M^3, \sigma_M^4, \sigma_C^1, \sigma_C^2, \sigma_C^3, \sigma_C^4$, ncpr, $\kappa >$ for a subset of samples in set 1 and samples generated by the model.** The model was trained on inputs whose $\kappa$ values were outside the range $[.4, .6]$. Points that are colored red refer to data from set 1 whose $\kappa$ value is within $[.4, .6]$, and points colored blue refer to data generated by the model. Note that to sample from the model, $\kappa$ was restricted between $[.4, .6]$. For reference, the number of instances in the actual and generated sample set is equal. Overall, we observe that the pairwise distributions of the generated samples strongly overlap with that of the actual samples. The x/y-axis of each plot stems corresponding column/row facet. Entries in the lower-triangular portion of the matrix represent scatter plots for each pair of variables while entries in the upper-triangular portion of the matrix represent 2d density contour plots. Entries on the diagonal represent 1d density plots.

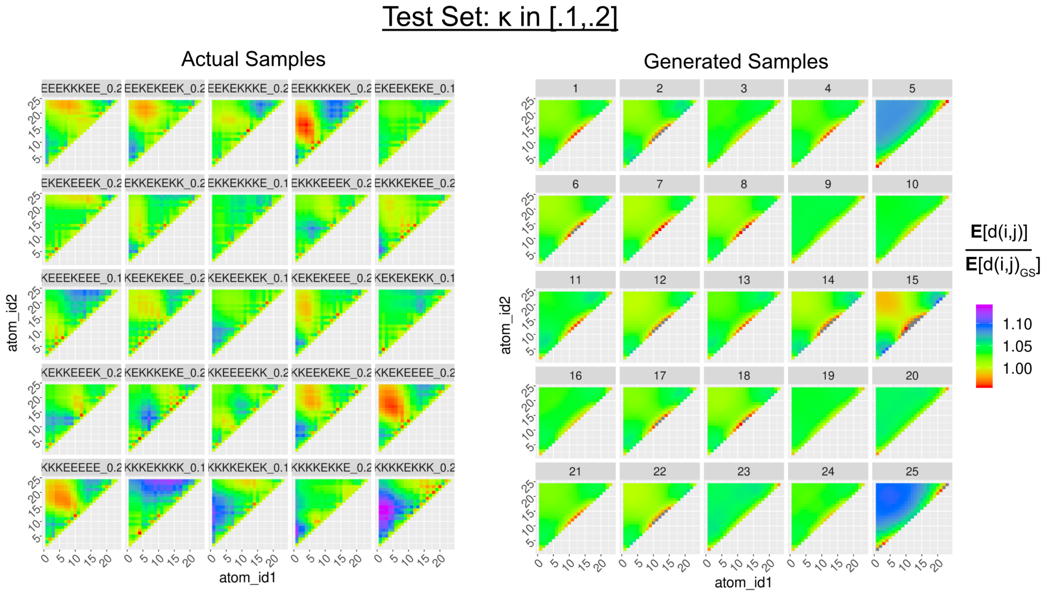


Figure S16. **Generation of averaged pairwise distance maps.** On the left, are actual samples from the $\kappa$ permutations set while on the right are samples generated by the model. The model was trained on inputs whose $\kappa$ values were outside the range $[.1, .2]$. Note that to sample from the model, $\kappa$ was restricted between $[.1, .2]$. In the distance map, each value represents the average distance between residue $i$ and $j$ divided by the average distance between residue $i$ and $j$ from the $(\mathrm{GS})_{12}$ simulation. Finally, we note that the color scale is based on the minimum and maximum value among the panel of plots of actual samples. To maintain ease of comparison, any value derived from the generated samples below or above the corresponding minimum or maximum value was set to gray.

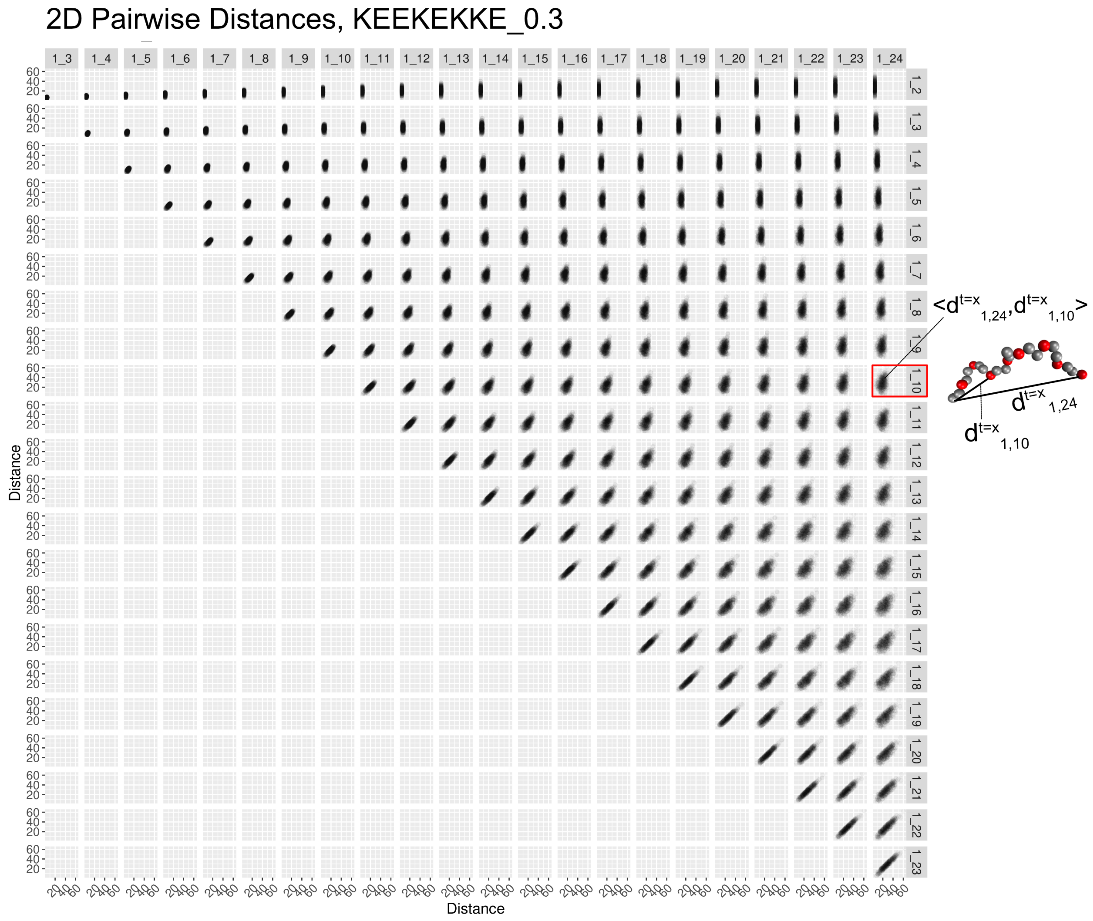


Figure S17. **Two-dimensional pairwise distance maps for a sequence from the $\kappa$ permutations set with $\kappa$ = .3.** Each subplot represents a sample whose x-coordinate represents the distance between residues $< i, j >$ and whose y-coordinate represents the distance between residues $< k, l >$ for a particular timeframe in the simulation. In this case, $i = k = 1$. In each subplot, we are displaying $< d_{ij}, d_{kl} >$ for every 100th frame, yielding 300 total points. For reference, the specific sequence corresponds to GSKKGSKKGSGSGSEGSEGSEGSE.

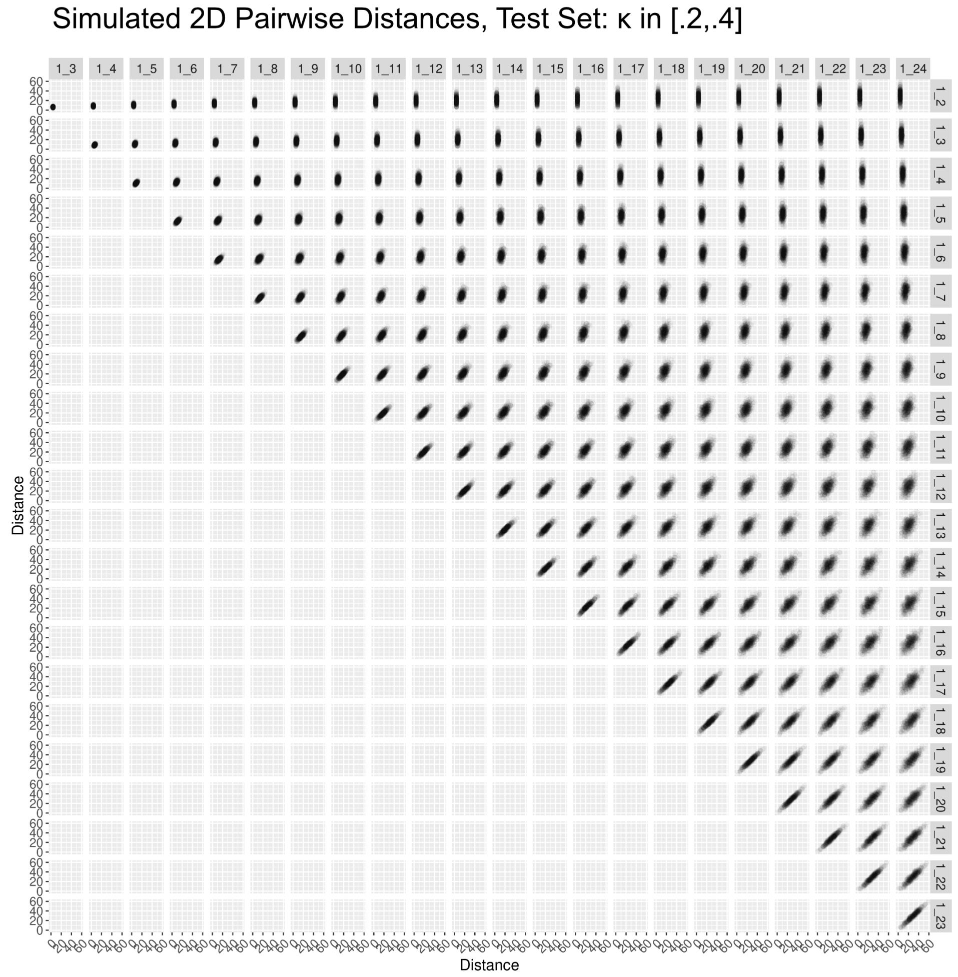


Figure S18. **Simulated two-dimensional pairwise distance maps. 300 samples are generated from a model whose test set consists of sequences whose $\kappa$ value is within** $[.2, .4]$**.** Each subplot represents a sample whose x-coordinate represents the distance between residues $< i, j >$ and whose y-coordinate represents the distance between residues $< k, l >$ for a particular timeframe in the simulation. In this case, $i = k = 1$.

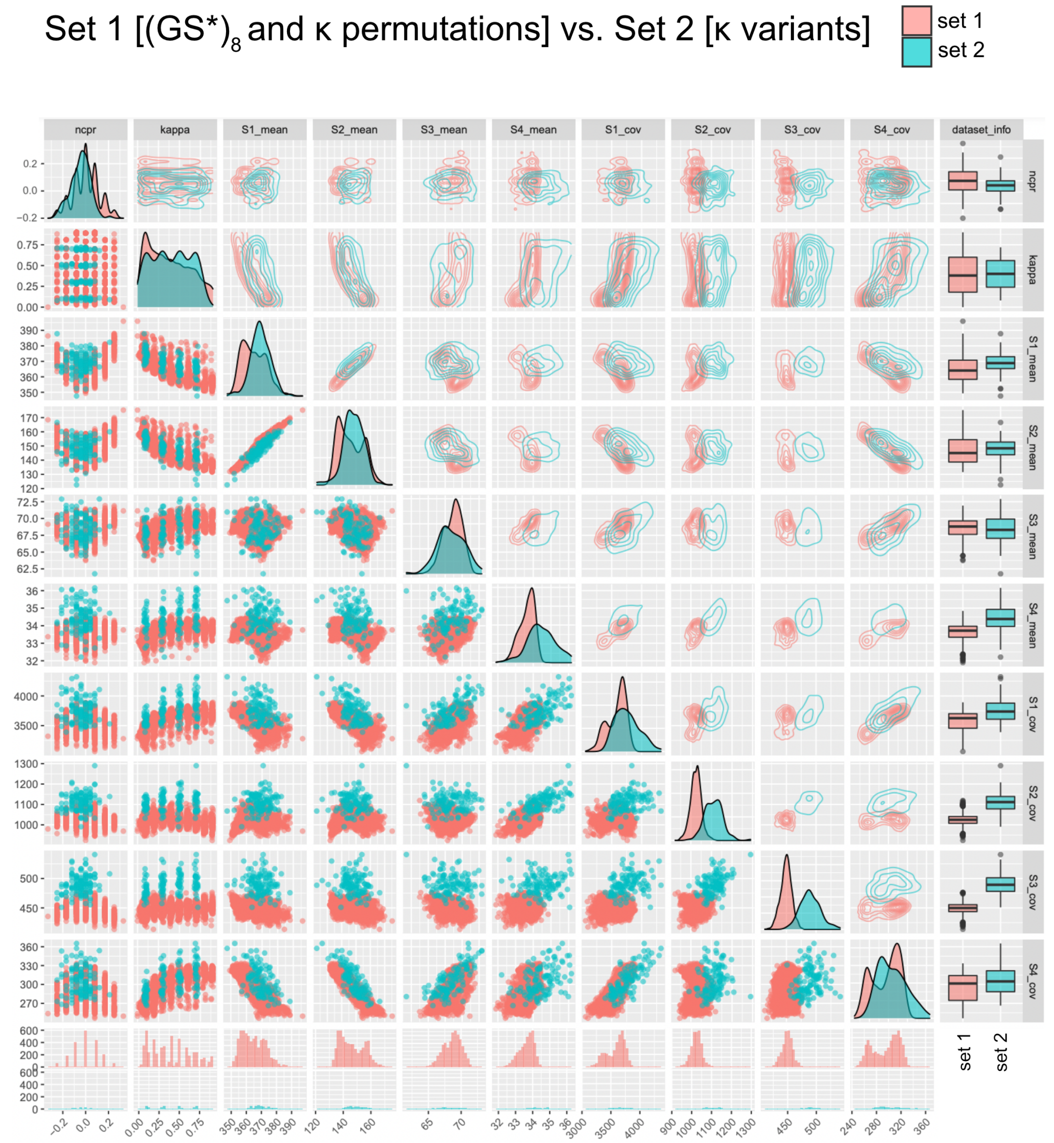


Figure S19. **Pairwise distribution for each pair of variables in $<\sigma_M^1, \sigma_M^2, \sigma_M^3, \sigma_M^4, \sigma_C^1, \sigma_C^2, \sigma_C^3, \sigma_C^4$, ncpr, $\kappa >$ for samples in set 1 and set 2.** Set 2 consists of all sequences from the $\kappa$ variants. Points that are colored red refer to data from set 1 (N = 1952) while points colored blue refer to data from set 2 (N = 120). The x/y-axis of each plot stems from the corresponding column/row facet. Entries in the lower-triangular portion of the matrix represent scatter plots for each pair of variables while entries in the upper-triangular portion of the matrix represent 2d density contour plots. Entries on the diagonal represent 1d density plots.

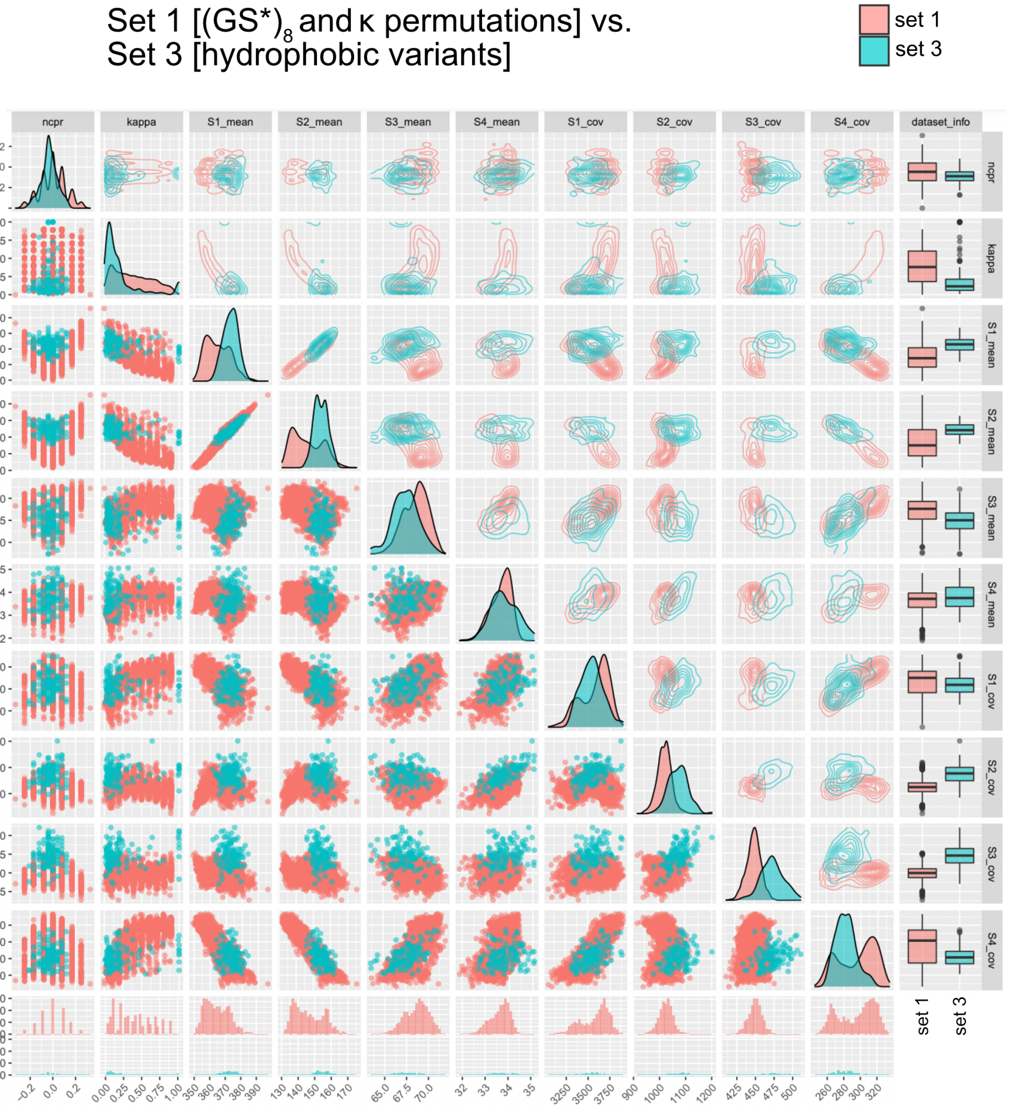


Figure S20. **Pairwise distribution for each pair of variables in $<\sigma_M^1, \sigma_M^2, \sigma_M^3, \sigma_M^4, \sigma_C^1, \sigma_C^2, \sigma_C^3, \sigma_C^4$, ncpr, $\kappa >$ for samples set 1 and set 3.** Set 3 consists of all sequences from the hydrophobic variants. Points that are colored red refer to data from set 1 (N = 1952) while points colored blue refer to data from set 3 (N = 120). The x/y-axis of each plot stems from the corresponding column/row facet. Entries in the lower-triangular portion of the matrix represent scatter plots for each pair of variables while entries in the upper-triangular portion of the matrix represent 2d density contour plots. Entries on the diagonal represent 1d density plots.

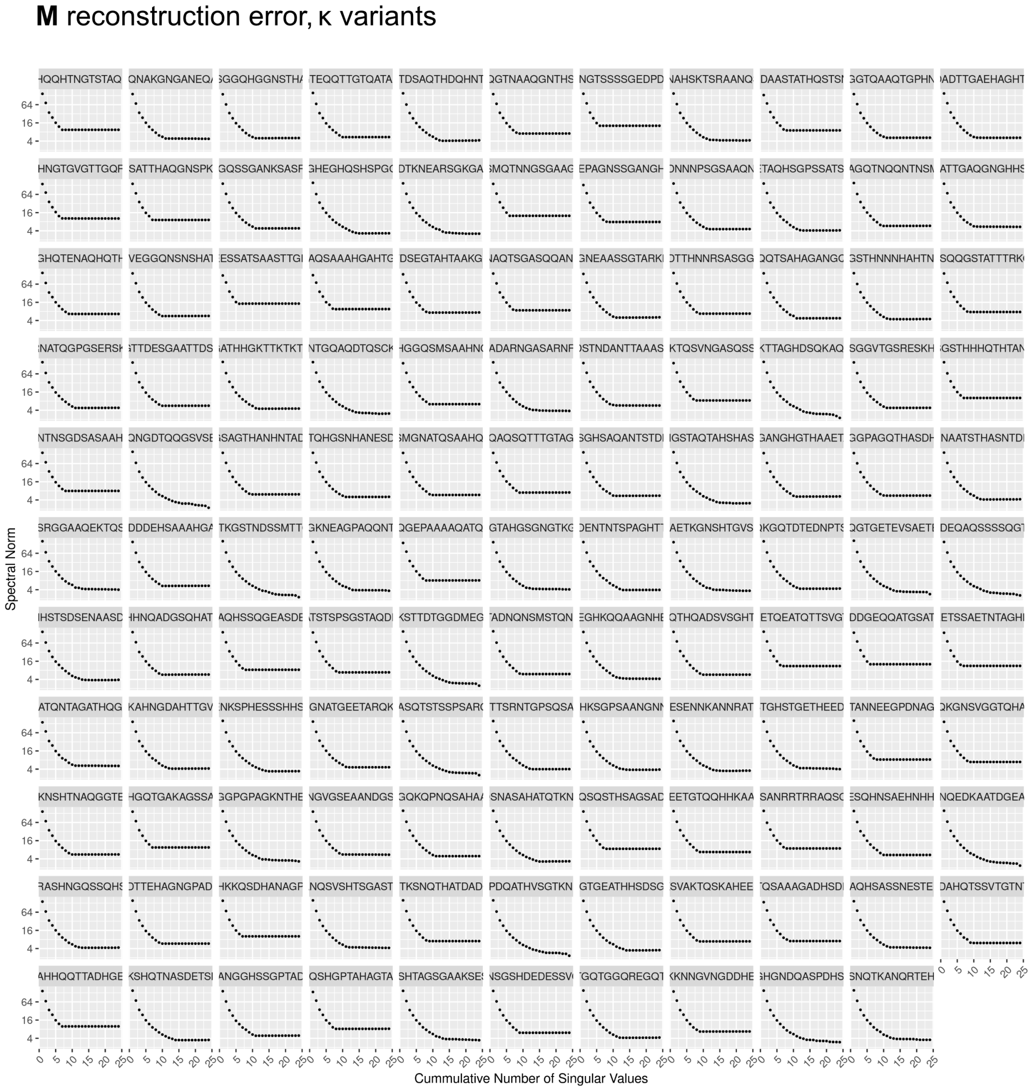


Figure S21. **Reconstruction error of M for each sequence in set 2 ($\kappa$ variants) as a function of the number of cumulative singular values used.** Reconstruction error is defined as the spectral norm between the difference of the reconstructed matrix and original matrix.

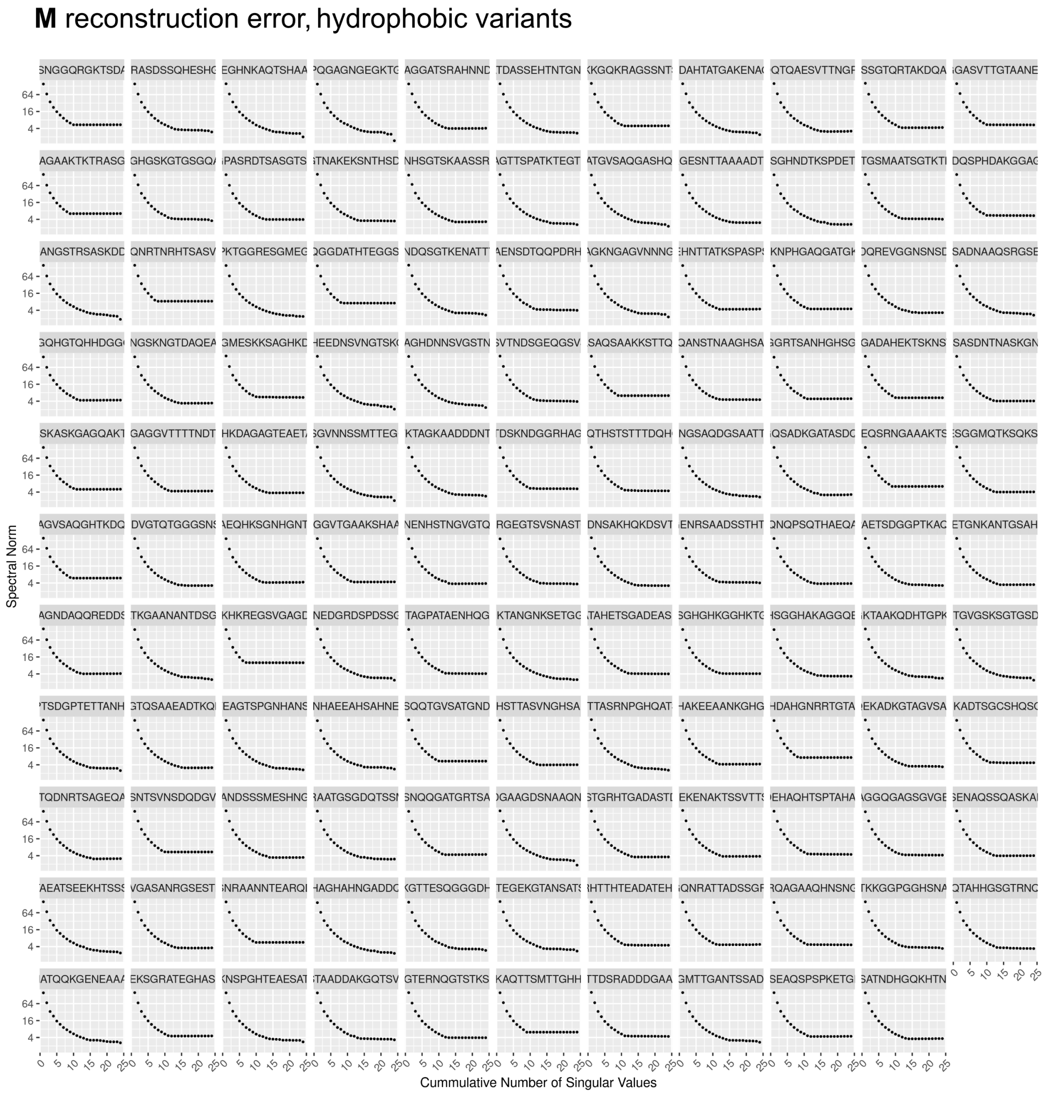


Figure S21. **Reconstruction error of M for each sequence in set 3 (hydrophobic variants) as a function of the number of cumulative singular values used.** Reconstruction error is defined as the spectral norm between the difference of the reconstructed matrix and original matrix.
